# Imperfect strategy transmission can reverse the role of population viscosity on the evolution of altruism

**DOI:** 10.1101/609818

**Authors:** F. Débarre

## Abstract

Population viscosity, *i.e.*, low emigration out of the natal deme, leads to high within-deme relatedness, which is beneficial to the evolution of altruistic behavior when social interactions take place among deme-mates. However, a detrimental side-effect of low emigration is the increase in competition among related individuals. The evolution of altruism depends on the balance between these opposite effects. This balance is already known to be affected by details of the life cycle; we show here that it further depends on the fidelity of strategy transmission from parents to their offspring. We consider different life cycles and identify thresholds of parent-offspring strategy transmission inaccuracy, above which higher emigration can increase the frequency of altruists maintained in the population. Predictions were first obtained analytically assuming weak selection and equal deme sizes, then confirmed with stochastic simulations relaxing these assumptions. Contrary to what happens with perfect strategy transmission from parent to offspring, our results show that higher emigration can be favorable to the evolution of altruism.

## Introduction

In his pioneering work on the evolution of social behavior, Hamilton suggested that al-truistic behavior would be associated to limited dispersal (Hamilton, 1964, p. 10). This notion, that tighter links between individuals are beneficial to the evolution of altruism, has been shown to hold in a number of population structures (see *e.g.* Ohtsuki et al., 2006; Taylor et al., 2007a; Lehmann et al., 2007; Allen et al., 2017). The rationale is that altruism is favored when altruists interact more with altruists than defectors do (Hamilton, 1975, p. 141; Fletcher & Doebeli, 2009), a condition that is met in viscous populations, i.e., populations with limited dispersal.

Yet, living next to your kin also implies competing against them (West et al., 2002; Platt & Bever, 2009), which is detrimental to the evolution of altruism. The evolution of social traits hence depends on the balance between the positive effects of interactions with related individuals and the detrimental consequences of kin competition. Under specific conditions, the two effects can even compensate each other, thereby annihilating the impact of population viscosity on the evolution of altruism. First identified with computer simulations (Wilson et al., 1992), this cancellation result was analyzed by Taylor (1992a) in a model with synchronous generations (*i.e.*, Wright-Fisher model) and a subdivided population of constant, infinite size. The cancellation result was later extended to heterogeneous populations (Rodrigues & Gardner, 2012, with synchronous generations and infinite population size), and other life cycles, with generic regular population structures (Taylor et al., 2011, with synchronous generations but also with continuous generations and Birth-Death updating). However, small changes in the model’s assumptions, such as overlapping generations (Taylor & Irwin, 2000) or the presence of empty sites (Alizon & Taylor, 2008) can tip the balance in the favor of altruism. This high dependence on life cycle specificities highlights the difficulty of making general statements about the role of spatial structure on the evolution of altruism.

Three different life cycles are classically used in studies on altruism in structured populations: Wright-Fisher, where the whole population is renewed at each time step, and two Moran life cycles (Birth-Death and Death-Birth), where a single individual dies and is replaced at each time step. We will consider the three of them in this study, because even though they differ by seemingly minor details, they are known to have very different outcomes in models with perfect parent-offspring transmission (*e.g.*, Taylor, 1992a; Rousset, 2004; Ohtsuki et al., 2006; Lehmann et al., 2007; Taylor, 2010).

A large number of studies on the evolution of social behavior consider simple population structures (typically, homogeneous populations *sensu* Taylor et al. (2007a)) and often also infinite population sizes (but see Allen et al., 2017, for results on any structure). These studies also make use of weak selection approximations, and commonly assume rare (*e.g.*, Leturque & Rousset, 2002; Taylor et al., 2007b; Tarnita & Taylor, 2014; Chen et al., 2019) or absent mutation (for models assuming infinite population sizes, or models concentrating on fixation probabilities; see Lehmann & Rousset, 2014; Van Cleve, 2015, for recent reviews). These simplifying assumptions are often a necessary step to-wards obtaining explicit analytical results. Simple population structures (*e.g.*, regular graphs, or subdivided populations with demes of equal sizes) help reduce the dimensionality of the system under study, in particular when the structure of the population displays symmetries such that all sites behave the same way in expectation. Weak selection approximations are crucial for disentangling spatial moments (Lion, 2016), that is, changes in global *vs.* local frequencies (though they can in some cases be relaxed, as in Mullon & Lehmann, 2014). Mutation, however, is usually ignored by classical models of inclusive fitness because these models assume infinite population sizes, so that there is no need to add mechanisms that restore genetic diversity (Tarnita & Taylor, 2014). In populations of finite size, this diversifying effect can be obtained thanks to mutation.

When strategy transmission is purely genetic, it makes sense to assume that mutation is relatively infrequent. Even in this case, though, mutations from “social” to “non-social” types cannot always be neglected. For instance, experiments with the bacteria *Pseudomonas fluorescens* have identified transitions between populations dominated by the ancestral “solitary” Smooth Morph type and mat-forming “social” Wrinkly Spreaders, that can be re-invaded by Smooth Morphs not contributing to the formation of the mat (hence described as “cheaters”). The transitions between the different types are due to spontaneous mutations occurring over the timescale of the experiment (Hammer-schmidt et al., 2014). In addition to genetic transmission, a social strategy can also be culturally transmitted from parent to offspring. In this case, “rebellion” (as in Frank’s Rebellious Child Model (Frank, 1997)), *i.e.*, adopting a social strategy different from one’s parents, does not have to be infrequent. Since it is known that imperfect strategy transmission can alter the evolutionary dynamics of social traits, in particular in spatially structured populations (see *e.g.*, Allen et al., 2012; Débarre, 2017, for graph-structured populations), it is therefore important to understand the impact of imperfect strategy transmission on the evolution of social behavior.

Here, we want to explore the consequences of imperfect strategy transmission from parents to their offspring on the evolution of altruistic behavior in subdivided populations^1^. The question was tackled by Frank (1997), but with a non “fully dynamic model” (Frank, 1997, legend of Fig.7). Relatedness was treated like a parameter, which precluded the exploration of the effects of population viscosity on the evolution altruism.

For each of the three life cycles that we consider, we compute the expected (*i.e.*, long-term) frequency of altruists maintained in a subdivided population, and investigate how this frequency is affected by mutation and emigration. We find that, contrary to what happens with perfect strategy transmission, higher emigration can increase the expected frequency of altruists in the population.

## Model and methods

### Assumptions

We consider a population of total size *N*, subdivided into *N*_*D*_ demes connected by dispersal, each deme hosting exactly *n* individuals (*i.e.*, each deme contains *n* sites, each of which is occupied by exactly one individual; *nN*_*D*_ *= N*). Each site has a unique label *i*, 1 ≤ *i* ≤ *N*. There are two types of individuals in the population, altruists and defectors. The type of the individual living at site *i* (1 ≤ *i* ≤ *N*) is given by an indicator variable *X*_*i*_, equal to 1 if the individual is an altruist, and to 0 if it is a defector. The state of the entire population is given by a vector **X** *=* {*X*_*i*_}_1≤*i* ≤*N*_. For a given population state **X**, the proportion of altruists is 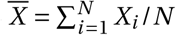. All symbols are summarized in table A1.

Reproduction is asexual. The offspring of altruists are altruists themselves with probability 1 −*µ*_1→0_, and are defectors otherwise (0 < *µ*_1→0_ ≤ 1/2). Similarly, the offspring of defectors are defectors with probability 1 −*µ*_0→1_, and are altruists otherwise (0 < *µ*_0→1_ ≤ 1/2). Our calculations will be simpler if we introduce the following change of parameters:

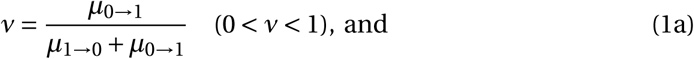

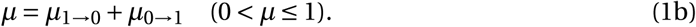

The composite parameter *v* corresponds to the expected frequency of altruists in the population at the mutation-drift balance (*i.e.*, in the absence of selection; see Appendix A for details). We call *ν* the “mutation bias” parameter. Parameter *µ* is the sum of the two mutation probabilities. In the absence of selection, at the mutation-drift equilibrium, the correlation between offspring type and their parent’s type is 1−*µ* (see Appendix A for details for the calculation). We call *µ* the mutation intensity.

An individual of type *X*_*k*_ expresses a social phenotype *φ*_*k*_ *= δX*_*k*_, where *δ* is assumed to be small (*δ* ≪ 1). This assumption of small phenotypic differences leads to weak selection. This type of weak selection is called “*δ*-weak selection” in Wild & Traulsen (2007). Social interactions take place within each deme; a focal individual interacts with its *n*− 1 other deme-mates. We assume that social interactions affect individual fecundity; *f*_*k*_ denotes the fecundity of the individual at site *k* (1 ≤ *k* ≤ *N*), which depends on deme composition. We denote by b the sum of the marginal effects of deme-mates’ phenotypes on the fecundity of a focal individual, and by −c the marginal effect of a focal individual’s phenotype on its own fecundity (c ≤ b; see system (A22) for formal definitions).

Offspring remain in the parental deme with probability 1 − *m* and land on any site of the parental deme with equal probability (including the very site of their parent). With probability *m*, offspring emigrate to a different deme, chosen uniformly at random among the *N*_*D*_ − 1 other demes. Denoting by *d*_*i j*_ the probability of moving from site *i* to site *j*, we have

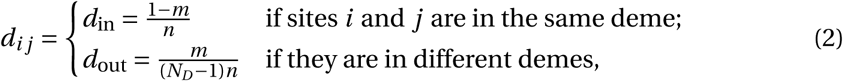

with 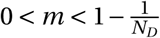. This upper bound is here to ensure that within-deme relatedness *R*, which will be defined later in the article, remains positive. When the emigration probability *m* is equal to the upper bound 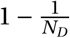, the population is effectively well-mixed (*d*_in_ *= d*_out_).

We denote by *B*_*i*_ *= B*_*i*_ (**X**, *δ*) the expected number of successful offspring of the individual living at site *i* (“successful” means alive at the next time step), and by *D*_*i*_ *= D*_*i*_ (**X**, *δ*) the probability that the individual living at site *i* dies. Both depend on the state of the population **X**, but also on the way the population is updated from one time step to the next, *i.e.*, on the chosen life cycle (also called updating rule). Because this term appears in our calculations, we also define

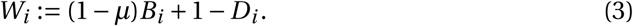

This is a particular definition of fitness, where the number of offspring produced (*B*_*i*_) is scaled by the parent-offspring type correlation (1 −*µ*).

We will specifically explore three different life cycles. At the beginning of each step of each life cycle, all individuals produce a large (effectively infinite) number of offspring, in proportion to their fecundity; some of these offspring can be mutated. Then these juveniles move, within the parental deme or outside of it, and land on a site. The next events occurring during the time step depend on the life cycle:

### Moran Birth-Death

One of the newly created juveniles is chosen at random; it kills the adult who was living at the site, and replaces it; all other juveniles die.

### Moran Death-Birth

One of the adults is chosen to die (uniformly at random among all adults). It is replaced by one of the juveniles who had landed in its site. All other juveniles die.

### Wright-Fisher

All the adults die. At each site of the entire population, one of the juveniles that landed there is chosen and establishes at the site.

Previous studies have shown that, when social interactions affect fecundity, altruism is disfavored under the Moran Birth-Death and Wright-Fisher life cycles, because the expected frequency of altruists under these life cycles is lower than what it would be in the absence of selection (*e.g.*, Taylor, 1992a, 2010; Taylor et al., 2011; Débarre, 2017). How-ever, we are interested in the actual value of the expected proportion of altruists in the population, not just whether it is higher or lower than the neutral expectation. This is why we are still considering the Moran Birth-Death and Wright-Fisher life cycles in this study.

## Methods

### Analytical part

The calculation steps to obtain the expected (*i.e.*, long-term) proportion of altruists are given in Appendix B. They go as follows: first, we write an equation for the expected frequency of altruists in the population at time *t* + 1, conditional on the composition of the population at time *t*; we then take the expectation of this quantity and consider large times *t*. After this, we write a first-order expansion for phenotypic differences *δ* close to 0 (this corresponds to a weak selection approximation).

The formula involves quantities that can be identified as neutral probabilities of identity by descent *Q*_*i j*_. These quantities correspond to the probability that individuals living at site *i* and *j* share a common ancestor and that no mutation occurred on either lineage since that ancestor, in a model with no selection (*δ =* 0) and with mutation intensity *µ*; this is the “mutation definition” of identity by descent (Rousset & Billiard, 2000). In a subdivided population like the one we consider, there are only three possible values of *Q*_*i j*_:

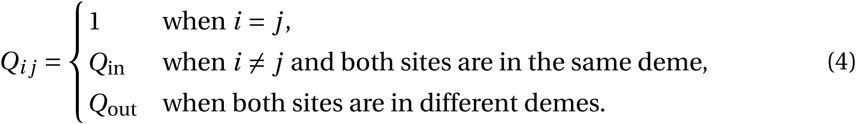

These neutral probabilities of identity by descent depend on the chosen life cycle, and are also computed by taking the long-term expectation of conditional expectations after one time step (see Appendix C.1 and C.2 and supplementary Mathematica file (Wolfram Research, Inc., 2017).)

### Stochastic simulations

To check our results and also relax some key assumptions, we ran stochastic simulations. The simulations were run for 10^8^ generations (one generation is one time step for the Wright-Fisher life cycle, and *N* time steps for the Moran life cycles). For each set of parameters and life cycle, we estimated the long-term frequency of altruists by sampling the population every 10^3^ generations and computing the average frequency of altruists.

All scripts are available at

https://flodebarre.github.io/SocEvolSubdivPop/

## Results

### Expected frequencies of altruists for each life cycle

For each of the life cycles that we consider, the expected frequency of altruists in the population,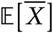, can be approximated as

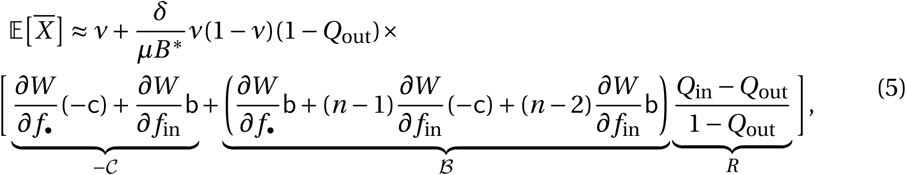

with *W* as defined in eq. (3). Calculations leading to eq. (5) are presented in Appendix B; notations are recapitulated in table A1. In particular, *B*\*is the expected number of off-spring produced by an adult, in the absence of selection (when *δ =* 0; *B*\**=* 1 for the Wright-Fisher life cycle and *B* * *=* 1/*N* for the Moran life cycles). Subscript “•” denotes a focal individual itself, and “in” a deme-mate. Partial derivatives are evaluated for *δ =* 0.

The expected frequency of altruists in the population is approximated, under weak selection (*δ* ≪ 1), by the sum of what it would be in the absence of selection 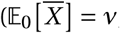, first term in eq. (5)), plus a deviation from this value, scaled by *δ*. The 𝒞 − term corresponds to the effects of a change of a focal individual’s phenotype on its own fitness (with the fitness definition given in eq. (3)). The ℬ term corresponds to the sum of the effects of the change of deme-mates’ phenotypes on an individual’s fitness. It is multiplied by *R*, which is relatedness.

The parametrization proposed in eq. (1) allows us to decouple the effects of the two new mutation parameters, *v* and *µ*. The mutation bias *v*, which was defined in eq. (1a), does not affect the sign of the second (“deviation”) term in eq. (5); it only appears in the *v*(1 − *v*) product. The mutation intensity *µ*, however, affects the values of *W*, *Q*_in_ and *Q*_out_. The presence of *µ* at the denominator in eq. (5) may look ominous; however, both *R* and (1 −*Q*_out_)/*µ* have a finite limit when *µ* → 0.

The different terms depend on the chosen life cycle. We first focus on relatedness *R*.

### Relatedness *R*

Within-deme relatedness *R* depends on the number of individuals that are born at each time step, and hence on the chosen life cycle. In a Moran life cycle (denoted by M), one individual is updated at each time step, while under a Wright-Fisher life cycle (denoted by WF), *N* individuals – the whole population – are updated at each time step. The formulas for relatedness, *R*^M^ and *R*^WF^, calculated for any number of demes *N*_*D*_ and mutation intensity *µ*, are presented in Appendix C.2 (eq. (A44) and eq. (A50)). When we let the number of demes go to infinity (*N*_*D*_ → *∞*) and the intensity of mutation be vanishingly small (*µ* → 0), we recover the classical formulas for relatedness as limit cases (eq. (A45) and eq. (A51)).

The effects of emigration *m* and mutation intensity *µ* on relatedness are represented in figure 1. For 0 < *m* < 1 − 1/*N*_*D*_, within-deme relatedness is positive, and it decreases with *m* and with *µ* (the mutation bias *v* has no effect). The effect of the mutation intensity *µ* on relatedness is strongest at low emigration probabilities *m*. As *m* increases, the relatedness values for different mutation intensities get closer, until they all hit zero for *m =* 1 − 1/*N*_*D*_ (which is the upper bound for the emigration values that we consider, a value such that there is no proper population subdivision anymore).

**Figure 1:**
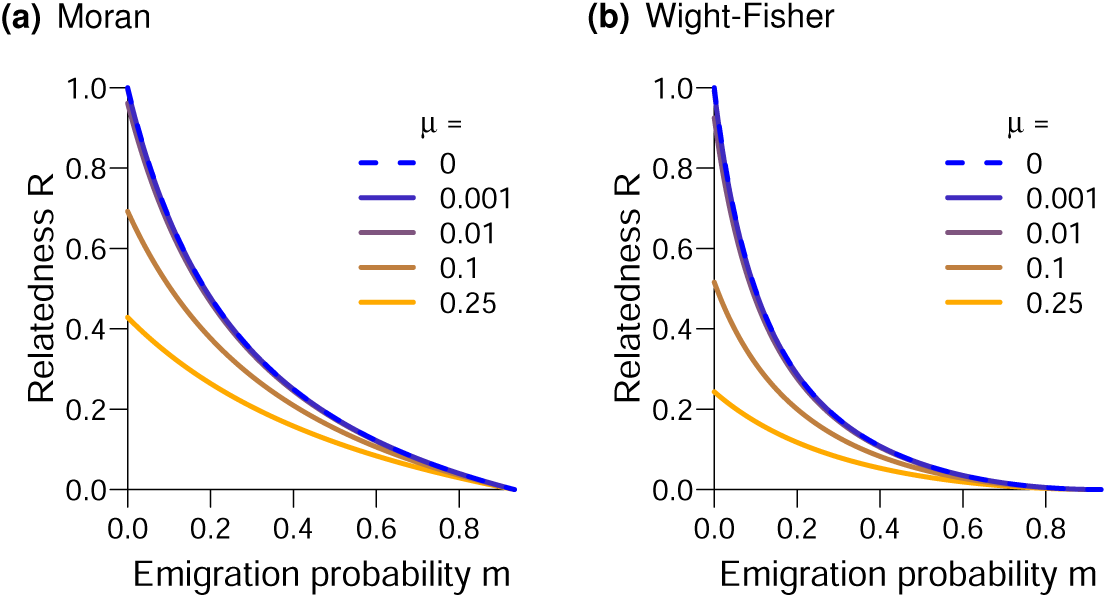
Within-deme relatedness of pairs of individuals *R*, as a function of the emigration probability *m*, for different values of the mutation probability *µ* (from 0 [blue] to 0.25 [orange]), and for the two types of life-cycles ((a): Moran, (b): Wright-Fisher). Other parameters: *n =* 4 individuals per deme, *N*_*D*_ *=* 15 demes.

### Primary and secondary effects

We now turn to the ℬ and − 𝒞 terms of eq. (5), which also depend on the chosen life cycle. We further decompose these terms into primary (subscript P) and secondary (subscript S) effects (West & Gardner, 2010):

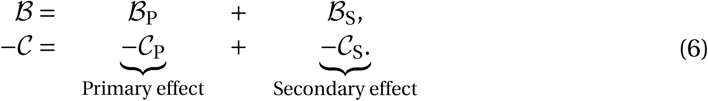

Primary effects correspond to unmediated consequences of interactions (they are included in 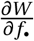). Secondary effects correspond to consequences of interactions mediated by other individuals, including competition.

### Primary effects

Primary effects are the same for all the life cycles that we consider:

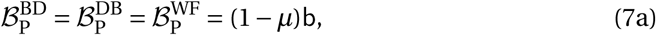

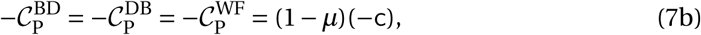

and they do not depend on the emigration probability *m* (see Appendix B.2 for details of the calculations).

As we have seen above, the relatedness terms *R*^M^ and *R*^WF^ decrease with *m* (keeping *m* < 1 − 1/*N*_*D*_; see figure 1). Consequently, if we ignored secondary effects, we would conclude that the expected frequency of altruists in the population 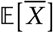 decreases as the emigration probability *m* increases. However, secondary effects play a role as well.

### Secondary effects

Secondary effects take competition into account, that is, how the change in the fecundity of an individual affects the fitness of another one. As shown already in models with nearly perfect strategy transmission (Grafen & Archetti, 2008), competition terms depend on the chosen life cycle, because life cycle details affect the distance at which competitive effects are felt. Given the way the model is formulated, − *𝒞*_S_ *= ℬ*_S_/(*n* − 1) holds for all the life cycles that we consider (see Appendix B.2 for details of the calculations).

Under the Moran Birth-Death life cycle, both the probability of reproducing and the probability of dying depend on the composition of the population. We obtain the following secondary effects:

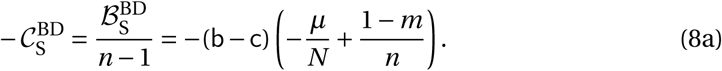

The competitive effects are the same for the Moran Death-Birth and Wright-Fisher life cycles. In both cases, the probabilities of dying are constant, so we can factor (1 −*µ*) in the equations:

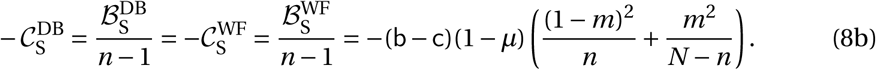

These secondary effects (eq. (8a) and eq. (8b)) remain negative for the range of emigration values that we consider (0 < *m* < 1 − 1/*N*_*D*_), and increase with *m*. In other words, the intensity of competition decreases as emigration *m* increases.

While the value of these secondary effects increases with emigration *m*, relatedness *R*, by which they are eventually multiplied in eq. (5), decreases with *m*. We therefore cannot determine the overall effect of emigration *m* on the expected frequency of altruists in the population by inspecting the different terms of eq. (5) in isolation. For each life cycle, we need to consider the entire equations to know the overall effect of the emigration probability *m* on the expected frequency of altruists 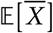 and on how it is affected by the (in)fideliy of parent-offspring transmission *µ*.

### Changes of the expected frequency of altruists with the emigration probability *m*

The rather lengthy formulas that we obtain are relegated to the Appendix and supplementary Mathematica file, and we concentrate here on the results.

### Moran Birth-Death

For the Moran Birth-Death life cycle, we find that the expected frequency of altruists 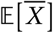 is a monotonic function of the emigration probability *m*. The direction of the change depends on the value of the mutation probability *µ* compared to a threshold value 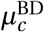. When 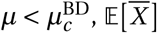 decreases with *m*, while when 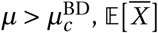 increases with *m*. The critical value 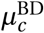 is given by

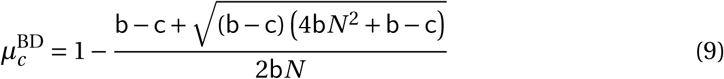

(recall that *N* is the total size of the population, *N = nN*_*D*_.) This result is illustrated in figure 2 (b); with the parameters of the figure, 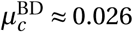. The threshold value increases with both deme size *n* and number of demes *N*_*D*_, up to a maximum value 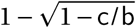 (equal to 0.034 with the parameters of figure 2(b).)

**Figure 2:**
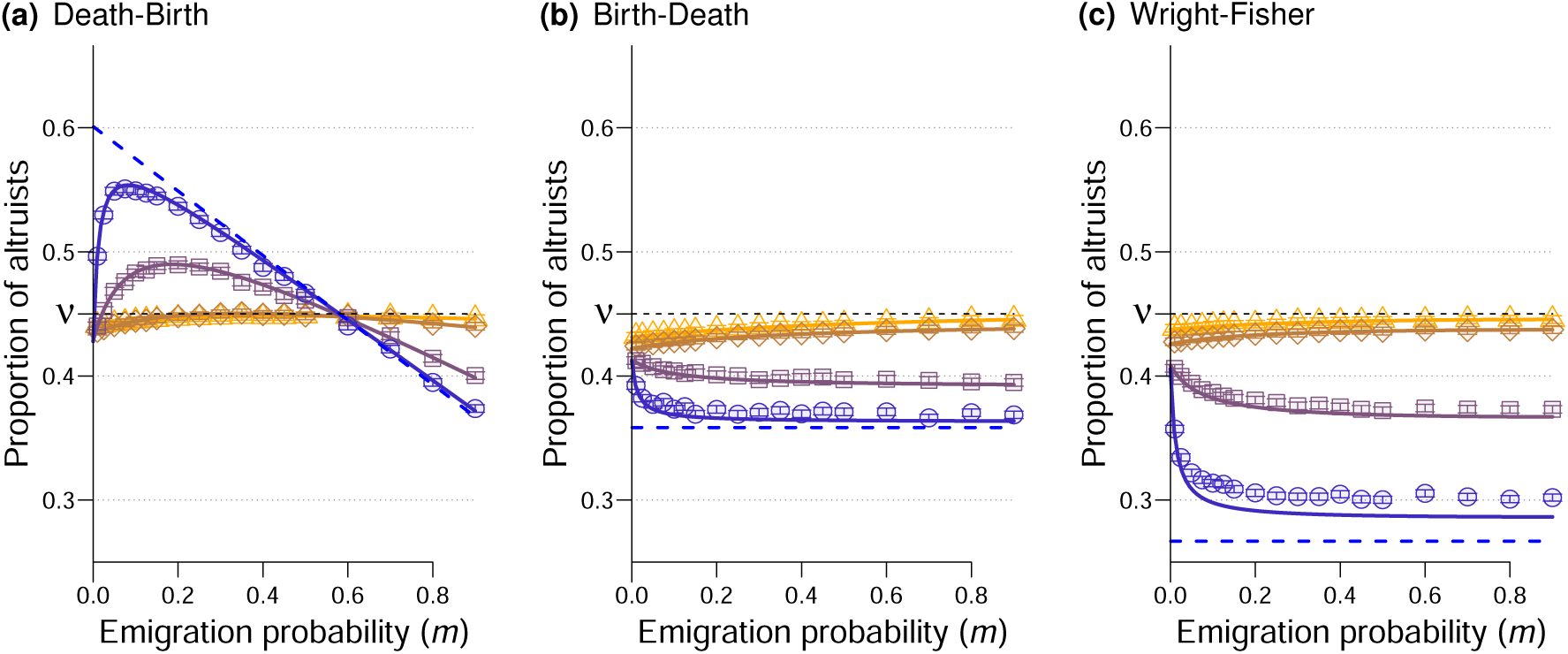
Expected proportion of altruists under weak selection, as a function of the emigration probability *m*, for different mutation values (*µ =* 0.001 (blue, dots), 0.01 (purple, squares), 0.1 (brown, diamonds), 0.25 (orange, triangles); the dashed blue lines correspond to *µ =* 0) and different life-cycles ((a) Moran Death-Birth, (b) Moran Birth Death, (c) Wright-Fisher). The curves are the analytical results, the points are the output of numerical simulations. Parameters: *δ =* 0.005, *ν =* 0.45, *b =* 15, *c =* 1, *n =* 4 individuals per deme, *N*_*D*_ *=* 15 demes.

With this life cycle however, the expected frequency of altruists 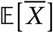 remains lower than *ν*, its value in the absence of selection (*i.e.*, when *δ =* 0).

### Moran Death-Birth

The relationship between 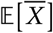 and *m* is a bit more complicated for the Moran Death-Birth life cycle. For simplicity, we concentrate on what happens starting from low emigration probabilities (*i.e.*, on the sign of the slope of 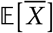 as a function of *m* when *m* → 0). If the benefits b provided by altruists are relatively low 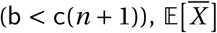 initially increases with *m* provided the mutation probability *µ* is greater than a threshold value *µ*^DB^ given in eq. (10) below; otherwise, when the benefits are high enough, 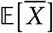 initially increases with *m* for any value of *µ*. Combining these results, we write

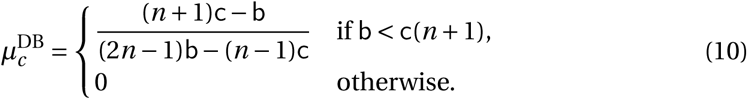

When b < c(*n*+1), the mutation threshold does not depend on the number of demes *N*_*D*_, but increases with deme size *n*. In figure 2(a), the parameters are such that 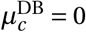.

When 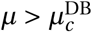, the expected frequency of altruists 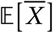 reaches a maximum at an emigration probability 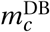 (whose complicated equation is given in the supplementary Mathematica file), as can be seen in figure 2 (a). When the mutation probability gets close to 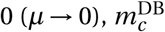 also gets close to 0.

With the Death-Birth life cycle, the expected frequency of altruists is higher than its neutral value ν for intermediate values of the emigration probability *m* (unless *µ* → 0, in which case the lower bound tends to 0).

### Wright-Fisher

Under a Wright-Fisher updating, the expected frequency of altruists in the population reaches an extremum at the highest admissible emigration value 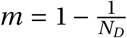. This extremum is a maximum when the mutation probability is higher than a threshold value 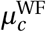 given by

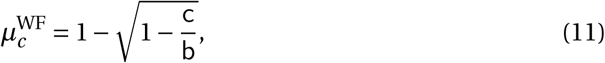

and it is a minimum otherwise. With the parameters of figure 2(c), 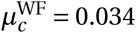.

With the Wright-Fisher life cycle however, the expected frequency of altruists remains below its value in the absence of selection, *ν*.

### Relaxing key assumptions

To derive our analytical results, we had to make a number of simplifying assumptions, such as the fact that selection is weak (*δ* ≪ 1), and the fact that the structure of the population is regular (all demes have the same size *n*). We checked with numerical simulations the robustness of our results when these key assumptions are relaxed.

### Strong selection

When selection is strong, the patterns that we identified not only still hold but are even more marked, as shown on figure A1.

### Heterogeneity in deme sizes

To relax the assumption of equal deme sizes, we randomly drew deme sizes at the beginning of simulations, with sizes ranging from 2 to 6 individuals and on average 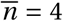 individuals per deme as previously. As shown in figure A2, the patterns initially obtained with a homogeneous population structure are robust when the structure is heterogeneous.

### No self-replacement

For the Moran model, it may seem odd that an offspring can replace its own parent (which can occur since *d*_*i i*_ ≠ 0). figure A3, plotted with dispersal probabilities preventing immediate replacement of one’s own parent (for all sites *i*, *d*_*i i*_ *= d*_self_ *=* 0; *d*_in_ *=* (1 −*m*)/(*n* − 1) for two different sites in the same deme, *d*_out_ remaining unchanged), confirms that this does affect our conclusions.

### Infinite number of demes

Our results are obtained in a population of finite size (the figures are drawn with *N*_*D*_ *=* 15 demes), but still hold when the size of the population is larger. Figure 3 (b) shows the range of emigration and mutation values such that altruism is favored, plotted also for *N*_*D*_ → ∞.

**Figure 3:**
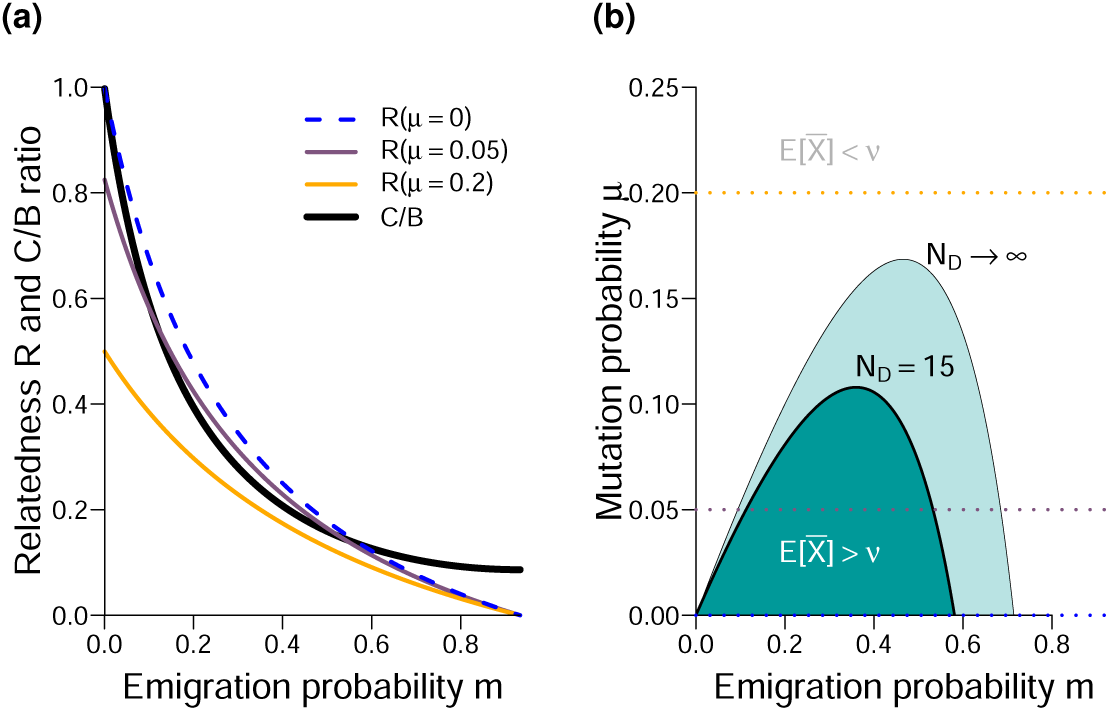
Understanding the effect of emigration *m* on whether altruism is favored in the Death-Birth life-cycle. (a) Comparison of the 𝒞/ℬ ratio (thick black curve) and relatedness *R* (thin curves) for different values of the mu<u>ta</u>tion probability *µ* (same color code as previously). (b) (*m, µ*) combinations for which 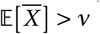. The dotted horizontal lines correspond to the mutation values used in panel (a). Unless specified, all other parameters are the same as in figure 2.

### Same graphs for dispersal and social interactions

Compared to graphs classically used in evolutionary graph theory (*e.g.*, regular random graphs, grids), the island model is particular because the interaction graph and the dispersal graph are different: interactions take place only within demes (*e*_out_ *=* 0), while offspring can disperse out of their natal deme (*d*_out_ > 0). One may wonder whether our result depends on this difference between the two graphs. figure A4 shows that the result still holds when the dispersal and interaction graphs are the same. In this figure indeed, we let a proportion *m* (equal to the dispersal probability) of interactions occur outside of the deme where the individuals live, and set *d*_self_, the probability of self replacement, equal to 0, so that the dispersal and interactions graphs are the same. Our conclusions remain unchanged.

## Discussion

### The expected frequency of altruists in a subdivided population can increase with the probability of emigration

Assuming that the transmission of a social strategy (being an altruist or a defector) from a parent to its offspring could be imperfect, we found that the expected frequency of altruists maintained in a population could increase with the probability *m* of emigration out of the parental deme, a parameter tuning population viscosity. This result can seem surprising, because it contradicts the conclusions obtained under the assumption of nearly perfect strategy transmission (*i.e.*, in the case of genetic transmission, when mutation is very weak or absent). Under nearly perfect strategy transmission indeed, increased population viscosity (*i.e.*, decreased emigration probability) is either neutral (Taylor, 1992a, and dashed lines in figures 2 (b)–(c)) or favorable (Taylor et al., 2007a, and dashed lines in figure 2 (a)) to the evolution of altruistic behavior.

### Quantitative vs. qualitative measures

Often, evolutionary success is measured qualitatively, by comparing a quantity (an expected frequency, or, in models with no mutation, a probability of fixation) to the value it would have in the absence of selection. In our model, this amounts to saying that altruism is favored whenever 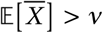 (*ν* is plotted as a horizontal dashed line in figure 2). Some of our conclusions change if we use this qualitative measure of evolutionary success: Under the Moran Birth-Death and Wright-Fisher life cycles, population viscosity does not promote the evolution of altruism – actually, these two life cycles cannot ever promote altruistic behavior for any regular population structure (Taylor et al., 2011), whichever the probability of mutation (Débarre, 2017). However, under a Moran Death-Birth life cycle (figure 2(a)), altruism can be favored only at intermediate emigration probabilities. Starting for initially low values of *m*, increasing the emigration probability can still favor the evolution of altruism under this qualitative criterion (see figure 3(b).)

### Interpreting the effect of *m* on 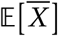

To better understand the role played by the mutation intensity *µ*, we focus on the qualitative condition for the evolution of altruism 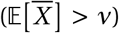; and on the Death-Birth life cycle, since this qualitative condition is not satisfied in the two other life cycles. Having made sure that ℬ ^DB^ > 0 (as shown in the supplementary Mathematical file), the qualitative condition for altruism to be favored is given by

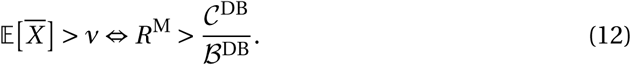

With the Death-Birth life cycle, the *C*^DB^/*B*^DB^ ratio does not change with the mutation probability *µ* (the (1 − *µ*) factors simplified out), but the ratio decreases with the emigration probability *m* (with 0 < *m* <1; − 1/*N*_*D*_; see the thick black curve in figure 3 (a)). This decrease of the 𝒞^DB^/ℬ^DB^ ratio is due to secondary effects (competition) diminishing as emigration increases. Relatedness, on the other hand, decreases with both *µ* and *m* (see figure 3 (a)). We need to explain the effect of the emigration probability *m* on condition (12) for different values of mutation intensity *µ*.

When the emigration probability *m* is high, relatedness gets closer to zero for all values of mutation intensity *µ*, while the 𝒞^DB^/ℬ^DB^ remains positive; condition (12) is not satisfied. On the other hand, when the emigration probability *m* is vanishingly small, 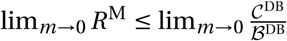, the two only being equal when *µ* = 0. Hence, condition (12) is satisfied for vanishingly low *m* only when strategy transmission is perfect. Finally, as *m* increases to intermediate values, the 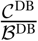 ratio decreases with a steeper slope than relatedness *R*, so that the curves can cross provided the mutation probability *µ* is not too high, *i.e.*, that *R* was not initially too low already. Hence, for no too high mutation intensity, there is a range of emigration values *m* such that condition (12) is satisfied.

### The result is due to secondary effects

The result, that frequency of altruists can increase with the emigration probability *m*, may seem counterintuitive. It is the case because verbal explanations for the evolution of altruism often rely on primary effects only. Relatedness *R* decreases with *m*, so it may be tempting to conclude that increases in the emigration probability *m* are necessarily detrimental to the evolution of altruism. However, secondary effects play an opposite role, as competition decreases with *m*, and the effect is strongest at low values of *m* (see the black curve on figure 3(a); in the absence of secondary effects, it would just be a horizontal line).

Secondary effects are less straightforward to understand than primary effects, and yet they play a crucial role for social evolution in spatially structured populations. Competition among relatives is for instance the reason for Taylor (1992b)’s cancellation result. Similarly, the qualitative differences between the Moran Birth-Death and Moran Death-Birth life cycles is explained by the different scales of competition that the two life cycle produce (Grafen & Archetti, 2008; Débarre et al., 2014). Secondary effects are also behind the evolution of social behaviors such as spite (West & Gardner, 2010).

### How small is small and how large is large?

Our results were derived under the assumption of weak selection, assuming that the phenotypic difference between altruists and defectors is small (*δ* ≪ 1). We considered any fidelity of transmission (any *µ* between 0 and 1) and population size. However, most models considering subdivided populations assume nearly perfect strategy transmission (*µ* → 0) and infinite population sizes (number of demes *N*_*D*_ → ∞). The point is technical, but it is important to know that the order in which these limits are taken matters, *i.e.*, one needs to specify how small *µ* and *δ* are compared to the inverse size of the population 1/*N*. This is in particular the case for the probability of identity by descent of two individuals in different demes, *Q*_out_: if we first take the small mutation limit, lim_*µ*→0_ *Q*_out_ *=* 0, while if we first take the large population limit, lim_*N*→∞_ *Q*_out_ *=* 1 (see Appendix C.2 for details). This remark complements findings by Sample & Allen (2017), who highlighted the quantitative differences between different orders of weak selection and large population limits.

### Imperfect transmission and Rebellious Children

Our model bears resemblance to the Rebellious Child Model by Frank (1997), who studied the evolution of a vertically transmitted cultural trait in an asexually reproducing population. In Frank’s model, however, relatedness *r* is treated as a fixed parameter (Frank, 1997, legend of Figure 7). Our model is mechanistic; relatedness *r* necessarily depends on the mutation probability *µ*, because probabilities of identity by descent do.

Mutation was also previously included in models investigating the maintenance of cooperative microorganisms in the presence of cheaters (Brockhurst et al., 2007; Frank, 2010). In both of these models however, only loss-of-function mutation was considered, which corresponds to setting the mutation bias at *ν =* 0 in our model. This means that the all-cheaters state is absorbing; no matter how favored cooperators may otherwise be, in the long run, a finite population will only consist of cheaters.

### Cultural transmission

Strategy transmission does not have to be genetic: it can be cultural. In our model, strategy transmission occurs upon reproduction, so this is a case of vertical cultural transmission.

The model could nevertheless be interpreted as a representation of horizontal transmission, if we described reproduction as an instance of an individual convincing another one to update its strategy. The Moran Death-Birth model can be interpreted as a modified imitation scheme (Boyd & Richerson, 2002; Ohtsuki et al., 2006; Traulsen et al., 2009)-with a specific function specifying who is imitated –, with mutation (Kandori et al., 1993), or as a voter model (Schneider et al., 2016). First, we choose uniformly at random an individual who may change its strategy; with probability *µ* the individual chooses a random strategy (altruistic with probability *ν*), and with probability 1 −*µ* it imitates another individual. Who is imitated depends on the distance to the focal individual (with probability *m* it is a random individual in another deme) and on the “fecundities” of those individuals (as shown in table A2). With this interpretation of the updating rule however, there is not reproduction nor death anymore.

It remains to be investigated how imperfect strategy transmission would affect the effect of population viscosity on the evolution of altruism in a model implementing both reproduction and horizontal cultural transmission (as in Lehmann et al., 2008). Such a model could then contrast the effects of impecfect genetic transmission and imperfect horizontal cultural transmission.

### Coevolution of dispersal and social behavior

This work also raises the question of what would happen if dispersal (*e.g.*, the emigration probability *m*) could evolve as well. Recent work on the topic has shown that under some conditions disruptive selection could take place, leading to a polymorphism between sessile altruists and mobile defectors (Parvinen, 2013; Mullon et al., 2017)— though more complex coevolutionary patterns can be obtained when considering the coevolution of altruism and mobility instead of natal dispersal, and unsaturated populations (Le Galliard et al., 2005). The assumptions of these studies however differ from ours in important ways, in that they consider continuous traits and use an adaptive dynamics framework, where, notably, mutations are assumed to be very rare. It remains to be investigated how non-rare and potentially large mutations would affect their result.

## Acknowledgements

Thanks to Charles Mullon and Jorge Peña for detailed comments on a previous version of the manuscript (CM suggested the *B R*−*C* decomposition; JP suggested the shortcut calculation of eq. (A17c)). Sébastien Lion suggested using the *R* vs. *𝒞* / *ℬ* comparison to interpret the result.

This work was funded by a ANR-14-ACHN-0003-01 grant.

## Supplementary figures

**Figure A1:**
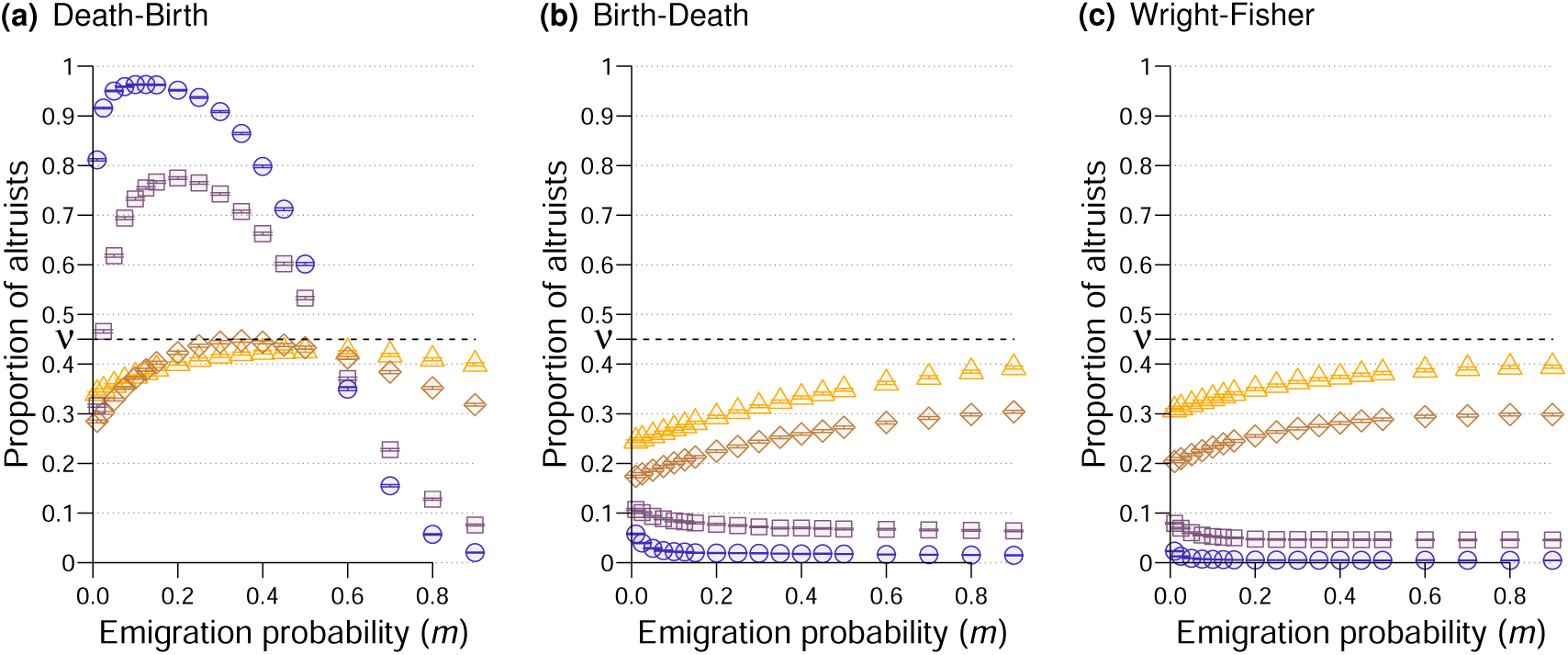
Equivalent of figure 2 (simulations only) but with strong selection (*δ =* 0.1); please note the change of scale on the vertical axis. All other parameters and legends are identical to those of figure 2 (increasing mutation probabilities from blue dots to orange triangles).

**Figure A2:**
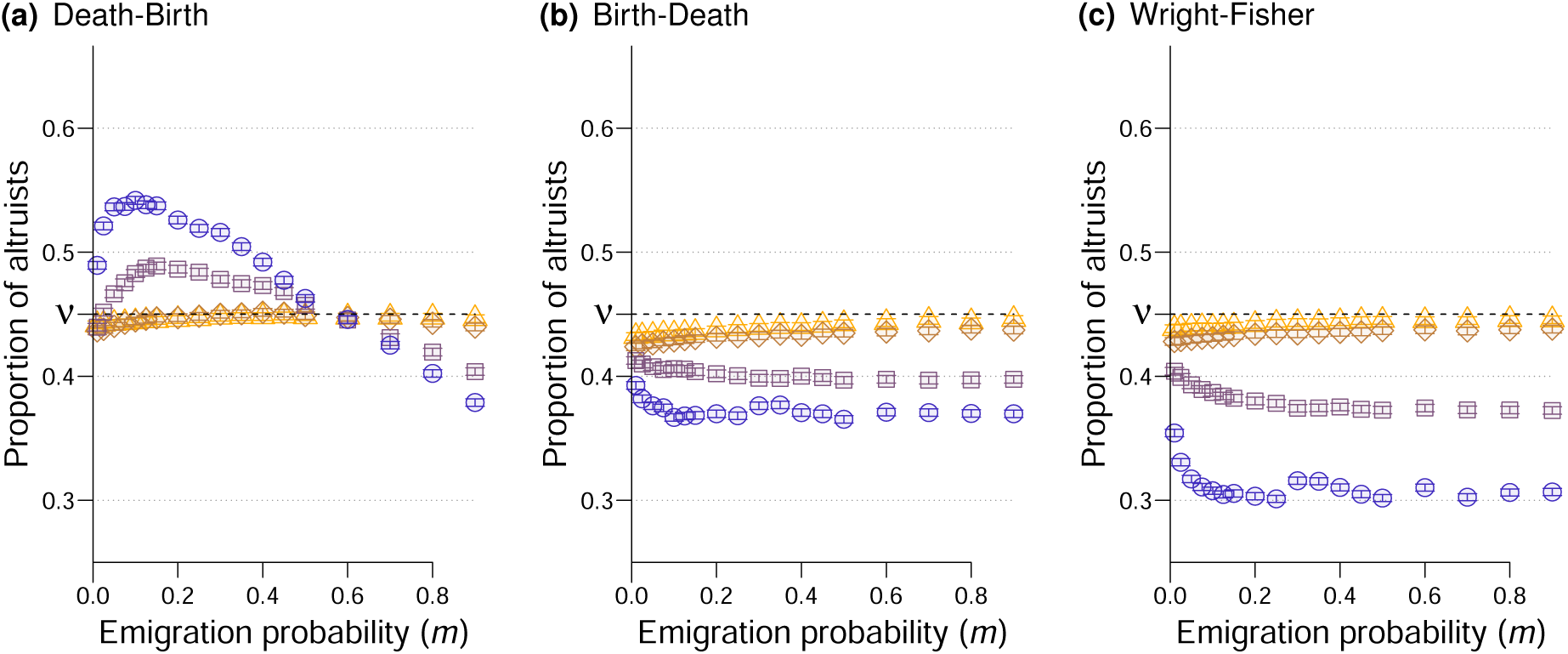
Equivalent of figure 2 (simulations only) but with a heterogeneous population structure: deme sizes range from 1 to 5 individuals per deme, the average deme size is 4 as in figure 2; all other parameters and legend are identical to those of figure 2.

**Figure A3:**
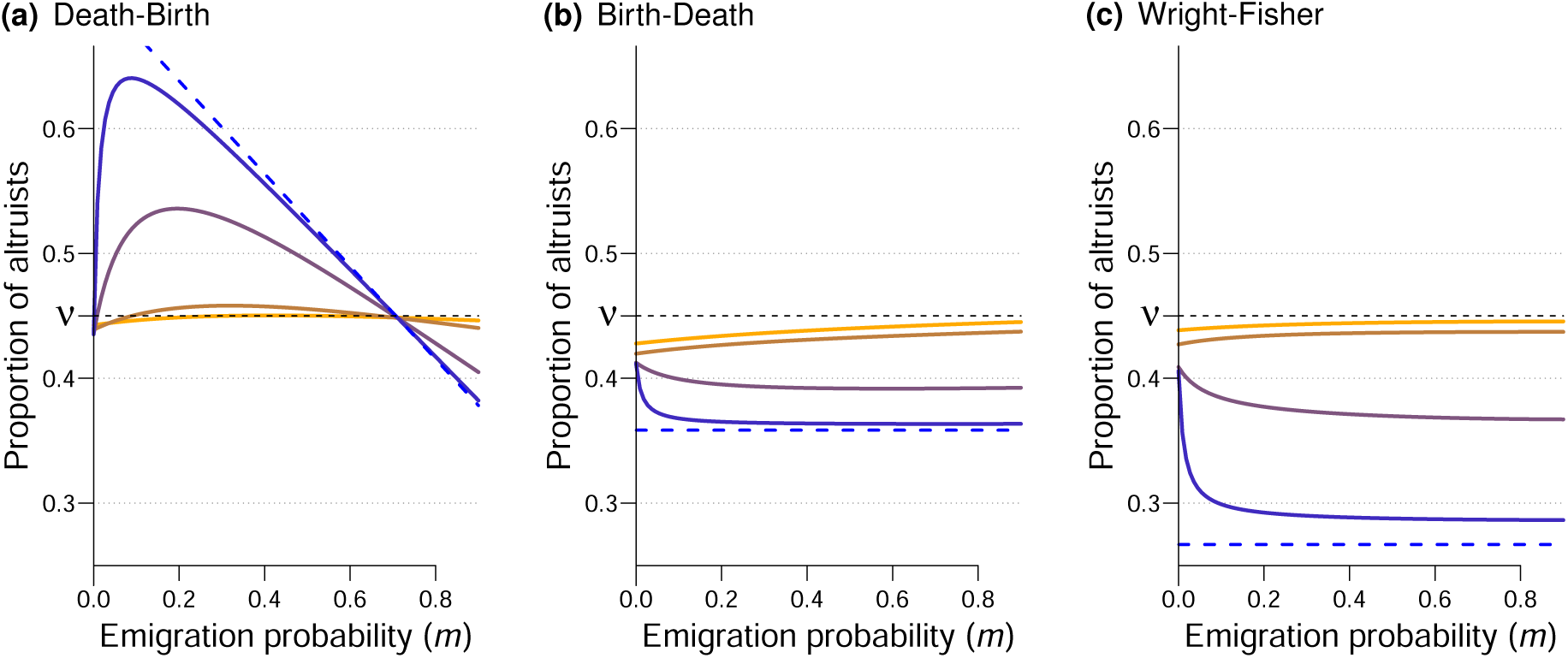
Equivalent of figure 2 (analysis only), with no self-replacement (*d*_*ii*_ *= d*_self_ = 0 for all sites).

**Figure A4:**
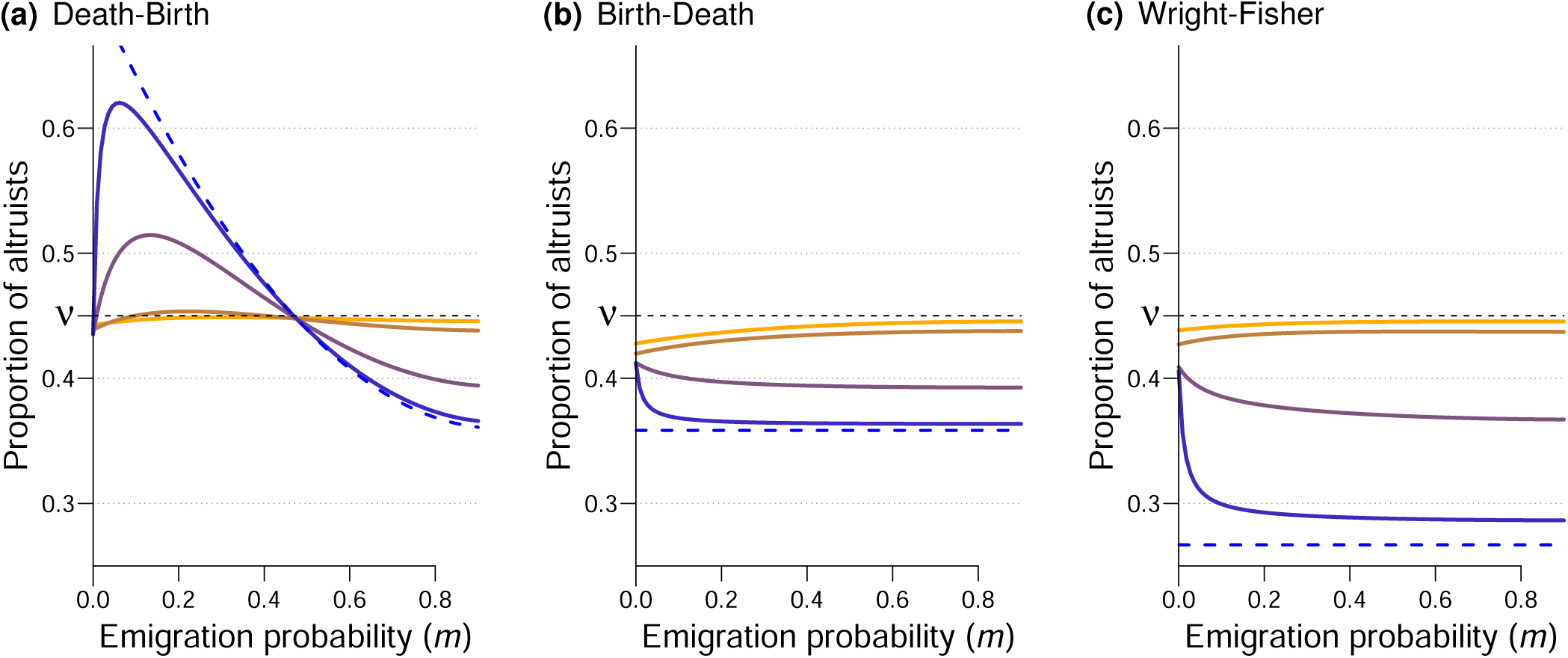
Equivalent of figure 2 (analysis only), with equal dispersal and interaction graphs (*i.e.*, no self-replacement [*d*_*ii*_ *= d*_self_ *=* 0 for all sites], and a proportion *m* of the interactions occurring outside of the home deme).

## Supplementary Table

**Table A1:** List of symbols. “r.v.” means *random variable*.

|  |  |
| --- | --- |
| $b$ | Sum of the marginal effects of deme-mates' phenotypes on focal individual's fecundity (benefit) |
| $\mathcal{B}$ | Sum of the marginal effects of deme-mates' phenotypes on the fitness $W$ of a focal individual |
| $B_i$ | Expected number of successful offspring of the individual living at site $i$ (r.v.) |
| $B^*$ | Value of $B_i$ for all sites, in the absence of selection ($\delta = 0$) |
| $c$ | Marginal effect of a focal individual's phenotype on its own fecundity (cost) |
| $C$ | Marginal effect of an individual's phenotype on its own fitness $W$ |
| $d_{ij}$ | Dispersal probability from site $i$ to site $j$ |
| $D_i$ | Probability that the individual currently living at site $i$ is dead at the end of the time step (r.v.) |
| $e_{ij}$ | Interaction probability from site $i$ to site $j$ |
| $f_i$ | Fecundity of the individual currently living at site $i$ (r.v.) |
| $n$ | Deme size |
| $N_D$ | Number of demes |
| $N$ | Total population size ($N = N_D n$) |
| $m$ | Emigration probability |
| $P_{ij}$ | (Long-term) Expected state of the pair of sites ($i, j$) |
| $Q_{ij}$ | (Long-term) Probability of identity by descent of individuals at sites $i$ and $j$ |
| $R$ | Pairwise within-deme relatedness (see eq. (5)) |
| $W_i$ | Measure of fitness, counting offspring only when unmutated (see eq. (3)) |
| $X_i$ | Indicator variable, equal to 1 if site $i$ is occupied by an altruist, to 0 otherwise (r.v.) |
| $\bar{X}$ | Frequency of altruists in the population (r.v.) |
| $\delta$ | Phenotypic distance between altruists and defectors; strength of selection |
| $\phi_i$ | Phenotype of the individual living at site $i$; $\phi_i = \delta X_i$ (r.v.) |
| $\mu$ | Mutation probability |
| $\nu$ | Mutation bias: probability that mutant is altruist |
| P | Subscript corresponding to primary effects |
| S | Subscript corresponding to secondary effects |
| • | Subscript used to denote a focal individual |
| in | Subscript used when $i \neq j$ and the two sites are in the same deme |
| out | Subscript used when the two sites $i$ and $j$ are in different demes |
| self | Subscript used when $i = j$ |
| 0 | Sub- or superscript meaning that a quantity is evaluated at $\delta = 0$ |
| BD | Superscript corresponding to the Moran Birth-Death model |
| DB | Superscript corresponding to the Moran Death-Birth model |
| M | Superscript corresponding to a Moran model |
| WF | Superscript corresponding to the Wright-Fisher model |

## Appendix

### A Mutation parameters

In the main text, we first introduce effective mutation parameters: *µ*_1→0_, the probability that an altruist has defector offspring, and *µ*_0→1_, the probability that a defector has altruist offspring.

#### A.1 Expected frequency of altruists at the mutation-drift balance

We assume that there is no selection acting (*δ =* 0), but that there still are two types of individuals in the population.

Let *Y* be the type of a randomly chosen individual (*Y =* 1 if the individual is an altruist and *Y =* 0 if it is a defector) in the population, given a proportion *y* of altruists in the population. In expectation, we have

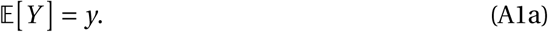

Let *Y* ′be the type of a randomly chosen individual at the next time step, given the frequency *y* at the previous time step. This randomly chosen individual is altruist if its parent was (which happens with probability *y*) and it did not mutate (probability 1 −*µ*_1→0_), or if its parent was not altruist (probability 1 − *y*), but the offspring mutated into one (probability *µ*_0→1_). We obtain

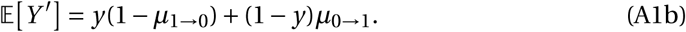

The expected frequency of altruists at the mutation-drift balance, denoted by *ν*, is found by solving 𝔼 [*Y*] *=* 𝔼[*Y*′]. We obtain

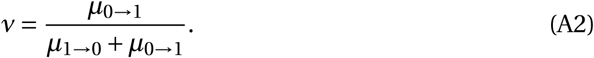

#### A.2 Parent-offspring correlation at the mutation drift balance

We can then compute the parent-offspring type correlation at the mutation-drift balance. First, let us compute the parent-offspring covariance:

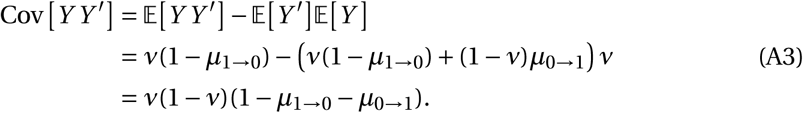

Remember that *Y* and *Y* ′are indicator variables and therefore take value in {0, 1}, so that *Y* ^2^ *= Y* (likewise for *Y* ′). Then, the standard deviations are given by

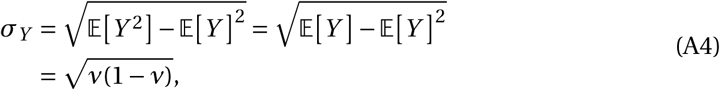

and

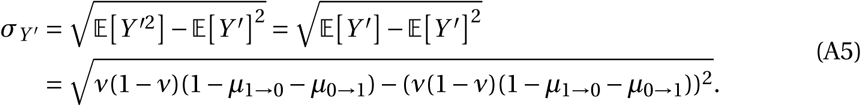

Finally, the parent-offspring correlation is given by

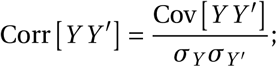

using the formulas eq. (A3)–(A5), and replacing *ν* by its value (mutation-drift equilibrium, eq. (A2)), we obtain

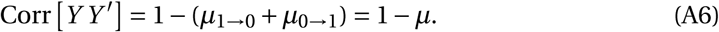

#### A.3 Redefining the mutation scheme

With the new mutation parameters *µ* and *ν*, we can describe the mutation scheme differently.

If we denote by *X*_*i*_ the type of a given parent, then the expected type of one of its offspring is

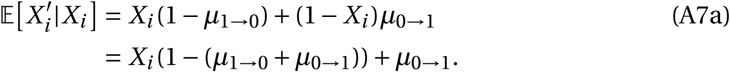

Replacing *µ*_1→0_ and *µ*_0→1_ by equivalent combinations of *µ* and *ν* as defined in eq. (A6) and eq. (A2), *i.e.*,

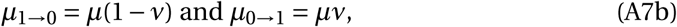

then eq. (A7a) becomes

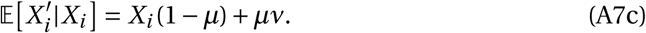

We can redefine the mutation scheme and interpret eq. (A7c) as follows. Parents transmit their strategy to their offspring with probability 1−*µ*; with probability *µ*, offspring do not inherit their strategy from their parent but instead get one randomly: with probability *ν*, they become altruists, with probability 1−*ν* they become defectors. With this alternative description, we can call “mutants” individuals who have the same type as their parent.

### B Expected frequency of altruists

#### B.1 For a generic life cycle

We want to compute the expected proportion of altruists in the population. We represent the state of the population at a given time *t* using indicator variables *X*_*i*_ (*t*), 1 ≤ *i* ≤ *N*, equal to 1 if the individual living at site *i* at time *t* is an altruist, and equal to 0 if it is a defector; these indicator variables are gathered in a *N*-long vector **X**(*t*). The set of all possible population states is Ω *=* {0, 1}^*N*^. The proportion of altruists in the population is written 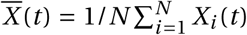. We denote by *B*_*j i*_ (**X**(*t*),*δ*), written *B*_*j i*_ for simplicity, the probability that the individual at site *j* at time *t* + 1 is the newly established offspring of the individual living at site *i* at time *t*. The expected number of successful offspring produced by the individual living at site *i* at time *t* is given by 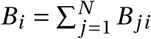. We denote by *D*_*i*_ (**X**(*t*), *δ*) (*D*_*i*_ for simplicity) the probability that the individual living at site *i* at time *t* has been replaced (*i.e.*, died) at time *t* + 1. These quantities depend on the chosen life cycle and on the state of the population; they are given in table A2 for each of the life cycles that we consider.

**Table A2:** Formulas of *B*_*ji*_ and *D*_*i*_ for each of the life cycles that we consider; *f*_*i*_ (shorthand notation for *f*_*i*_ (*X*, *δ*)) is the fecundity of the individual living at site *i*, and *d*_*ji*_ is a dispersal probability, given in eq. (2) in the main text.

| life cycle | $B_{ji}$ | $D_i$ |
| --- | --- | --- |
| Moran Birth-Death | $d_{ij} \frac{f_i}{\sum_{k=1}^N f_k}$ | $\frac{\sum_{j=1}^N d_{ji} f_j}{\sum_{k=1}^N f_k}$ |
| Moran Death-Birth | $\frac{1}{N} \frac{d_{ij} f_i}{\sum_{k=1}^N d_{kj} f_k}$ | $\frac{1}{N}$ |
| Wright-Fisher | $\frac{d_{ij} f_i}{\sum_{k=1}^N d_{kj} f_k}$ | 1 |

Since a dead individual is immediately replaced by one new individual (*i.e.*, population size remains constant and equal to *N*),

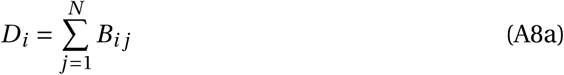

holds for all sites *i* and all life cycles.

The structure of the population is also such that in the absence of selection (*δ =* 0, so that *f*_*i*_ *=* 1 for all sites 1 ≤ *i* ≤ *N*), all individuals have the same probability of dying and the same probability of having successful offspring (*i.e.*, of having offspring that become adults at the next time step), so that

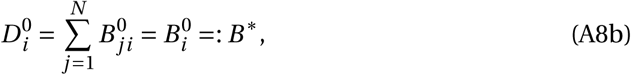

where the ^0^ subscript means that the quantities are evaluated for *δ =* 0. This also implies that 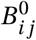 and 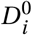 do not depend on the state **X** of the population. For the Moran life cycles, *B*\**=* 1/*N*, while for the Wright-Fisher life cycle, *B*\**=* 1. (The difference between eq. (A8b) and eq. (A8a) is that we are now considering offspring produced by *i* landing on *j*).

Given that the population is in state **X**(*t*) at time *t*, the expected frequency of altruists at time *t* + 1 is given by

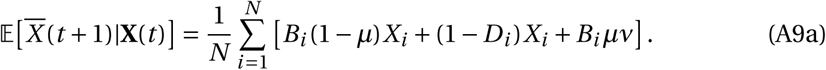

The first term within the brackets corresponds to births of unmutated offspring from parents who are altruists (*X*_*i*_). The second term corresponds to the survival of altruists. The third term corresponds to the births of mutants who became altruists (which occurs with probability *ν*), whichever the type of the parent.

A lost strategy can always be created again by mutation, so there is no absorbing population state. There exists a stationary distribution of population states (Theorem 1 in Allen & Tarnita (2014)). In other words, for large times *t*, the expected frequency of altruists does not change anymore (of course, realized frequencies keep changing over time). We denote by *ξ*(**X**, *δ, µ*) the probability that the population is in state **X**, given the strength of selection *δ* and the mutation probability *µ*. Taking the expectation of eq. (A9a) 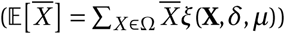, we obtain, after reorganizing:

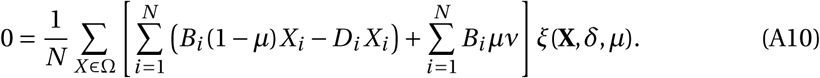

Now, we use the assumption of weak selection (*δ* ≪ 1) and consider the first-order expansion of eq. (A10) for *δ* close to 0.

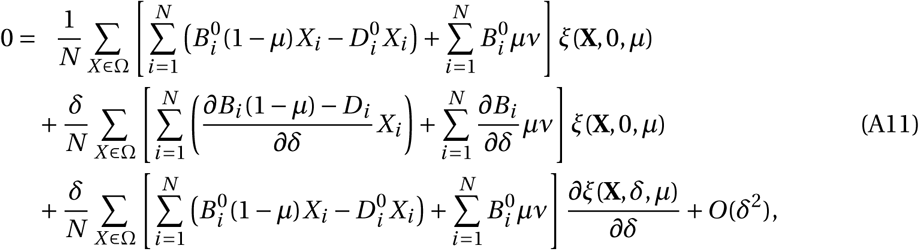

where all the derivatives are evaluated for *δ =* 0. The first line of eq. (A11) is equal to zero, because 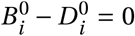 (eq. (A8b)), and because in the absence of selection (*δ =* 0), the expected state of every site *i* is 𝔼_0_ *X*_*i*_ *=* Σ_*X* ∈Ω_ *X*_*i*_ *ξ*(*X*, 0, *µ*) *= ν* (by definition of *ν*, see Appendix A.1). The second term of the second line is zero, because for all the life cycles that we consider, the total number of births in the population during one time step 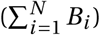 does not depend on population phenotypic composition (it is exactly 1 death for the Moran life cycles, and exactly *N* for the Wright-Fisher life cycle); since it is a constant, its derivative is 0. The third line simplifies by noting again that 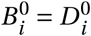 (first term), and that 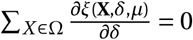 since *ξ* is a probability distribution (so the second term is zero). eq. (A11) then becomes

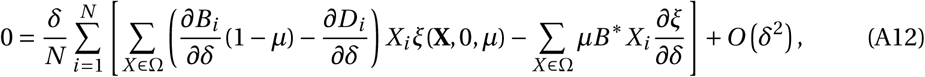

where the derivatives are evaluated at *δ =* 0. For conciseness, we define

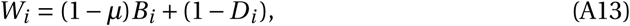

a measure of fitness counting offspring only when they are unmutated (in the sense of the alternate mutation scheme described in Appendix A.3). With this, using the expectation notation, and denoting by 𝔼_0_[] expectations under *δ =* 0, we can rewrite and reorganize eq. (A12) as

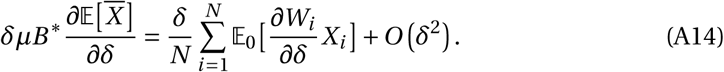

Now, we use a first time the law of total probabilities, taking individual phenotypes *ϕ*_*k*_ are intermediate variables:

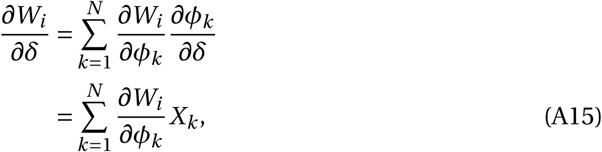

by definition of *ϕ*_*k*_ (*ϕ*_*k*_ *= δX*_*k*_), and where the derivatives are evaluated for all *ϕ*_*i*_ *=* 0, 1 ≤ *i* ≤ *N*. Introducing the notation *P*_*i j*_ *=* 𝔼_0_ [*X*_*i*_ *X* _*j*_] (expected state of a pair of sites), eq. (A14) becomes

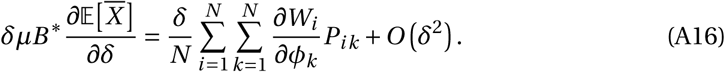

We note that *P*_*i i*_ *=* 𝔼_0_[*X*_*i*_ *X*_*i*_] *=* 𝔼_0_[*X*_*i*_] *= ν* (*X*_*i*_ being an indicator variable, it is either equal to 0 or 1, so 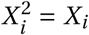). Given that the size of the population is fixed 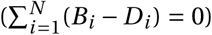, and given that the total number of births does not depend on population composition in the life cycles that we consider, we have

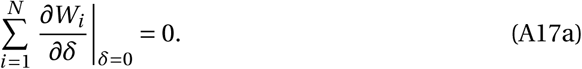

Using the decomposition in eq. (A15), which is valid for any population composition, and so in particular for **X** *=* **1**, eq. (A17a) becomes

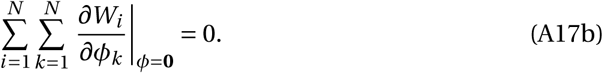

So far, we have not used the specificities of the population structure that we consider. First, the population is homogeneous (*sensu* Taylor et al., *2007a*). *Because this population homogeneity, eq. (A17b) is valid for all i* (not just their sum). Secondly, we are considering an island model. Once we have fixed a focal individual *i*, in expectation there are only three types of individuals: the focal itself (denoted by “•”), *n* − 1 other individuals in the focal’s deme (denoted by “in”), and *N* −*n* individuals in other demes (denoted by “out”). With these considerations, eq. (A17b) becomes

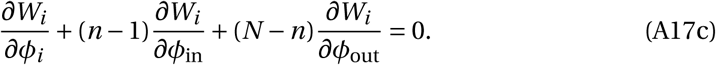

(as previously shown by (Rousset & Billiard, 2000, p.817–818)). Using this island model-specific notation, eq. (A16) becomes

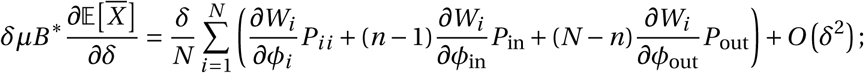

Injecting eq. (A17c) into eq. (A16), we obtain

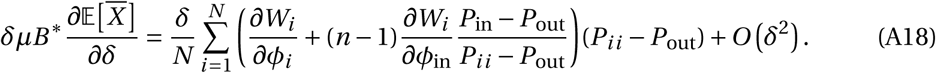

We can also replace the *P* terms as follows:

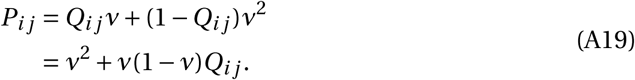

In Appendix C.1, using recursions on *P*_*i j*_, we will see that *Q*_*i j*_ can be interpreted as a probability of identity by descent, *i.e.*, the probability that the individuals at sites *i* and *j* have a common ancestor and that no mutation (using the alternative mutation scheme described in Appendix A.3) has occurred on either lineage since the ancestor. Replacing the *P* terms with eq. (A19), and noting that *Q*_*i i*_ *=* 1, eq. (A18) becomes

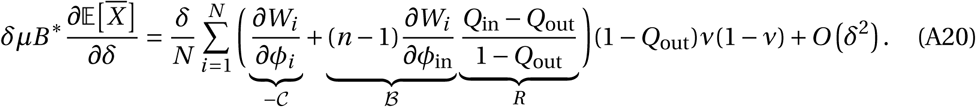

We can further decompose the derivatives, now using the fecundities *f*_*l*_ as intermediate variables, *i.e.*,

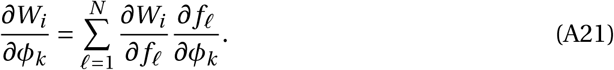

The term 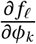 is the marginal effect of a change in the phenotype of the individual living at site *k* on the fecundity of the individual living at site 𝓁. By assumption, social interactions take place within demes only, so whenever sites 𝓁 and *k* are in different demes, we have 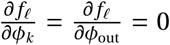. We then need to characterize the effect of one’s own phenotype (*i.e., k =* 𝓁) and of another deme-mate’s phenotype (*k* and 𝓁 being different sites in the same deme) on fecundity. For this, we define b and c so that:

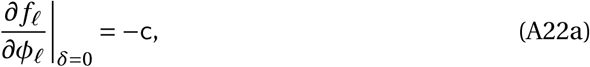

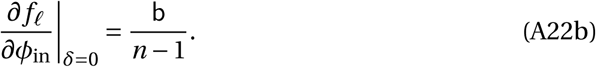

Eq. (A20) then becomes (using notation • to refer to the focal individual itself, and where *W = W*_*i*_, since the derivatives are the same for all *i*):

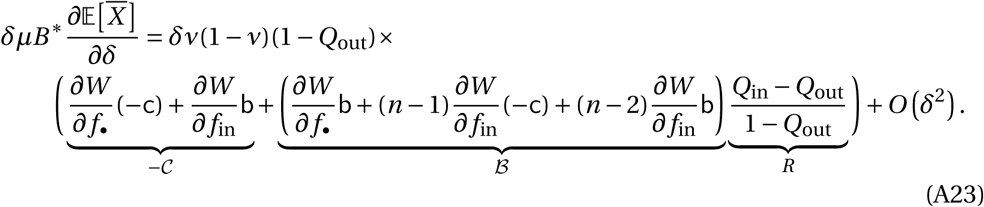

(As previously, all derivatives are evaluated at *δ =* 0.)

Finally, we write a first-order approximation of the expected frequency of altruists in the population:

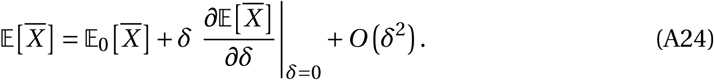

The first term, 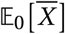, is the expected frequency in the absence of selection; it is equal to *ν* (as introduced in eq. (A2)). The derivative 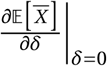 is obtained from eq. (A23). We then need to replace the *B*_*i*_ and *D*_*i*_ terms by their formulas for each life cycle; they are given in table A2. This is how the expected frequency of altruists in the population is approximated.

#### B.2 Derivatives for the specific life cycles

We use the formulas presented in table A2 and the definition of *W = W*_*i*_ given in eq. (A13) for each life cycle. In eq. (A26), eq. (A28) and eq. (A30), the first lines within parentheses correspond to primary effects, and the second line to secondary effects.

##### Moran Birth-Death

Under this life cycle, we obtain

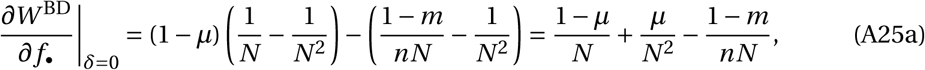

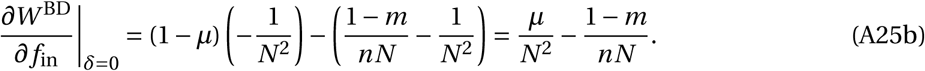

With these derivatives, eq. (5) becomes

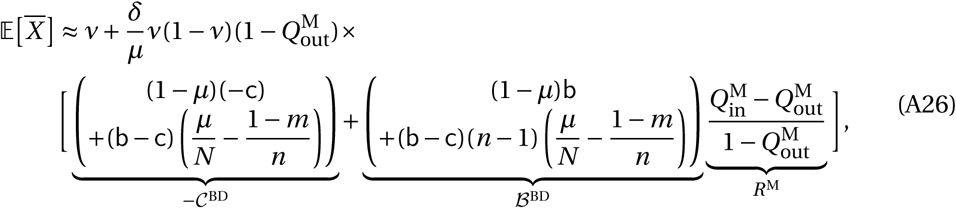

In addition, for both Moran life cycles, we have 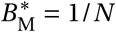. The secondary effects (second line in the parentheses in eq. (A26)) include competitive effects on the probability of reproducing, and consequences of social interactions on the probability that a given individual dies. Note that the secondary effects remain negative for the realistic range of emigration values that we consider (*i.e., m* < 1 − 1/*N*_*D*_).

##### Moran Death-Birth

Under this life cycle, we obtain

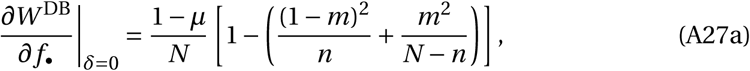

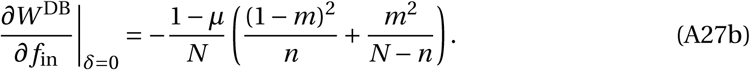

With the Death-Birth life cycle, eq. (5) becomes

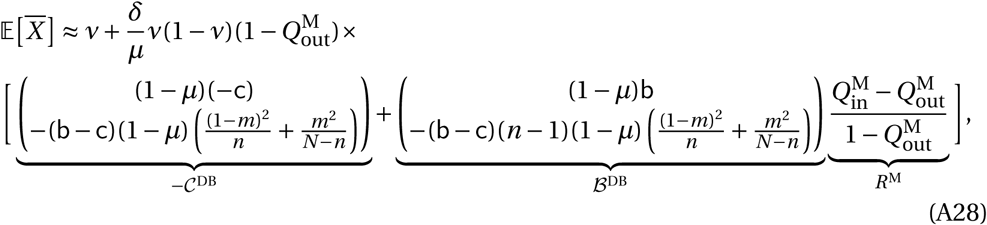

With this life cycle, Death occurs first, and the probability of dying is independent from the state of the population (since we assume that social interactions affect fecundity. We can therefore factor (1 − *µ*) in all terms. The primary effects (first lines in the parentheses) remain the same as with the Birth-Death life cycle. However, the Death-Birth life cycle leads to different secondary effects compared to the Birth-Death life cycle: competition occurs at a different scale (Grafen & Archetti, 2008). Finally, with this life cycle as we defined it, the probabilities of identity by descent *Q* are the same as with the Birth-Death model.

##### Wright-Fisher

Under this life cycle, we obtain

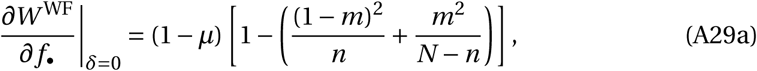

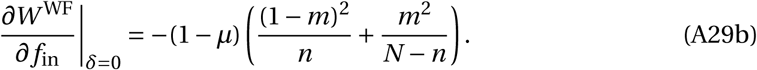

For the Wright-Fisher life cycle, we have 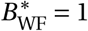. Replacing the derivatives presented in eq. (A29) into eq. (5), we obtain

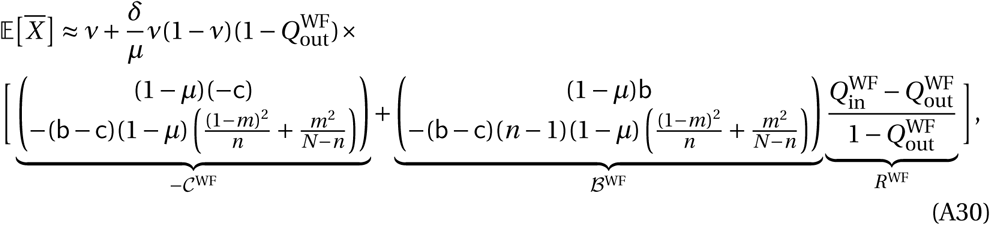

The only – but important – difference between eq. (A30) and eq. (A28) is the value of the probabilities of identity by descent *Q*, because the number of individuals that are updated at each time step differs.

### C Probabilities of identity by descent

#### C.1 Expected state of pairs of sites and probabilities of identity by descent

Here we show the link between the expected state of a pair of sites *P*_*i j*_ and probabilities of identity by descent *Q*_*i j*_. In our derivation of 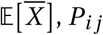 is the quantity that appears, but most studies use *Q*_*i j*_. Both are evaluated in the absence of selection (*δ =* 0).

#### C.1.1 Moran model

These calculations apply to both the Death-Birth and Birth-Death updating rules.

In a Moran model, exactly one individual dies and one individual reproduces during one time step. Given a state **X** at time *t*, at time *t* + 1 both sites *i* and *j* ≠ *i* are occupied by altruists, if *i*) it was the case at time *t* and neither site was replaced by a non-altruist (first term in eq. (A31)), or *ii*) if exactly one of the two sites was occupied by a non-altruist at time *t*, but the site was replaced by an altruist (second and third terms of eq. (A31)):

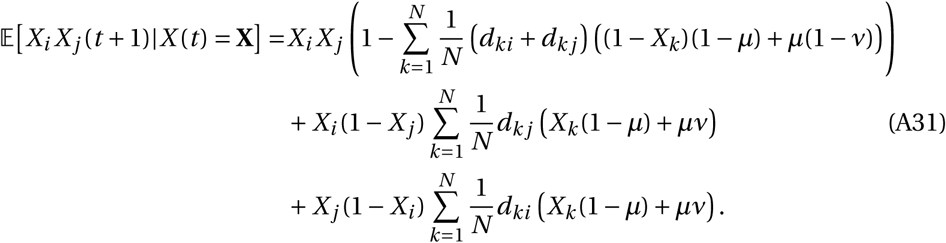

We take the expectation of this quantity, and consider that the stationary distribution is reached (*t* → ∞); then 𝔼[*X*_*i*_ *X* _*j*_ (*t* + 1)] *=* 𝔼[*X*_*i*_ *X* _*j*_ (*t*)], and we obtain after a few lines of algebra:

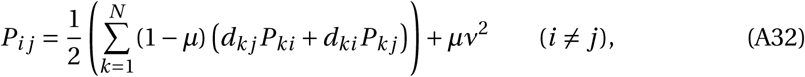

while *P*_*i i*_ *= ν*.

Now we substitute *P*_*i j*_ *= ν*^2^ + *ν*(1 −*ν*)*Q*_*i j*_ in eq. (A32), we obtain

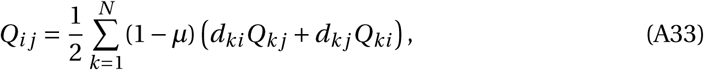

and we realize that *Q*_*i j*_ is the probability that the individuals at sites *i* and *j* ≠ *i* are identical by descent (*e.g.*, Taylor et al. (2011), equation above (S1.11); Allen & Nowak (2014) eq. (4)). To compute it indeed, we need to pick which site was last updated (*i* or *j* with equal probabilities: 1/2), then sum over the possible parent (*k*); the other individual needs to be identical by descent to the parent (*Q*_*k j*_, *Q*_*ki*_), disperse to the considered site (*d*_*ki*_, *d*_*k j*_), and no mutation should have occurred (1 −*µ*).

##### C.1.2 Wright-Fisher model

In a Wright-Fisher model, all individuals are replaced at each time step, so we directly consider the state of the parents:

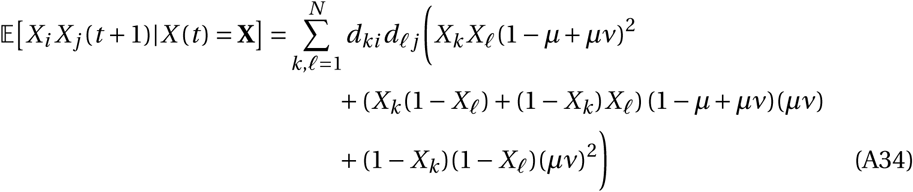

The first term of eq. (A34) corresponds to both parents being altruists, and having altruist offspring; the second line corresponds to exactly one parent being altruist, and the third line to both parents being non-altruists (in this latter case, the two offspring have to be both mutants to be altruists).

Taking the expectation and simplifying, we obtain

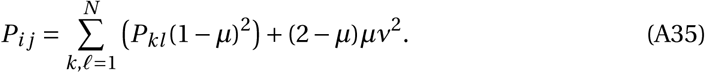

Replacing *P*_*i j*_ by ν^2^ + *ν*(1 −*v*)*Q*_*i j*_, eq. (A35) becomes

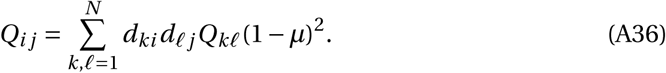

Again, *Q*_*i j*_ corresponds to a probability of identity by descent: the individuals at sites *i* and *j* are identical by descent if their parents were and if neither mutated ((1 −*µ*)^2^).

#### C.2 Probabilities of identity by descent in a subdivided population

Two individuals are said to be identical by descent if there has not been any mutation on either lineage since their common ancestor. Because of the structure of the population, there are only three types of pairs of individuals, and hence three different values of the probabilities of identity by descent of pairs of sites *Q*_*i j*_:

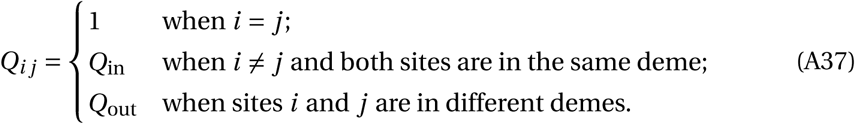

The values of *Q*_in_ and *Q*_out_ depend on the type of life cycle that we consider.

When the number of demes is infinite, *Q*_in_ is relatively easily obtained using recurrence equations and noting that *Q*_out_ *=* 0. However, writing the recurrence equations for *Q*_in_ and *Q*_out_ is much more tedious for finite populations. Hence, for finite populations, we will use formulas already derived in Débarre (2017) for “two-dimensional population structures”. The name comes from the fact that we only need two types of transformations to go from any site to any other site in the population: permutations on the deme index, and permutations on the within-deme index.

We rewrite site labels (1 ≤ *i* ≤ *N*) as (*ℓ*_1_, *ℓ*_2_), where *ℓ*_1_ is the index of the deme (1 ≤ *ℓ*_1_ ≤ *N*_*D*_) and *ℓ*_2_ the position of the site within the deme (1 ≤ *ℓ*_2_ ≤ *n*). Then, we introduce notations 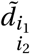 and 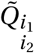, that correspond to the dispersal probability and probability of identity by descent to a site at distances *i*_1_ and *i*_2_ in the among-demes and within-deme dimensions 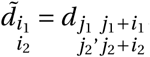

Also, in this section, we distinguish between *d*_self_ *= d*_*i i*_ and *d*_in_ (in the main text, *d*_self_ *= d*_in_).

##### C.2.1 Moran model

In Débarre (2017), it was shown that

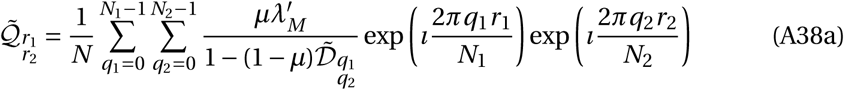

With

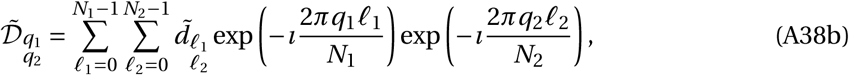

and 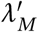 such that 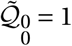. Let us first compute 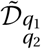 in the case of a subdivided population, with *N*_1_ *= N*_*D*_ and *N*_2_ *= n*:

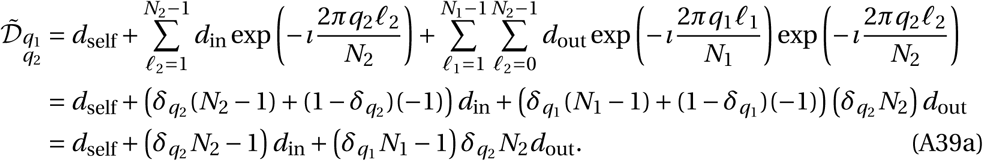

(*δ*_*q*_ is equal to 1 when *q* is equal to 0 modulo the relevant dimension, and to 0 otherwise). So for the three types of distances that we need to consider (distance 0, distance to another deme-mate, distance to individual in another deme), and with *N*_1_ *= N*_*D*_ and *N*_2_ *= n*, we obtain

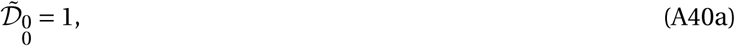

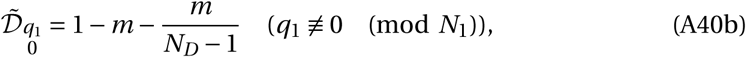

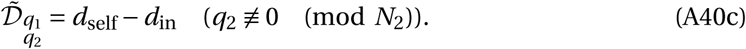

So for 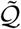, using system (A40) in eq. (A38a),

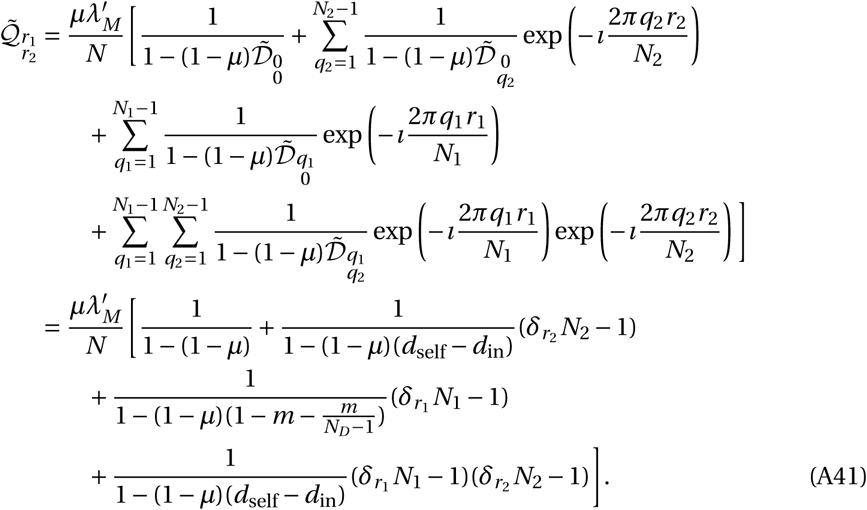

In particular,

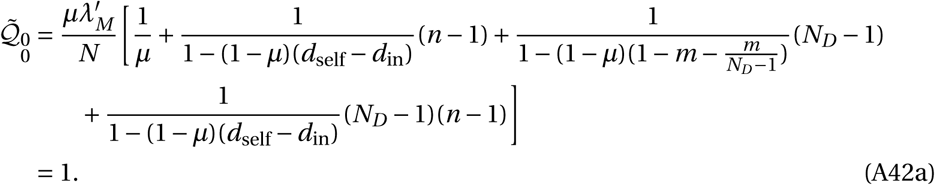

We find 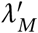 using eq. (A42a). Let’s now go back to eq. (A41): when *r*_1_ *=* 0, the two individuals are in the same deme. The two individuals are different when *r*_2_ ≢ 0, and so:

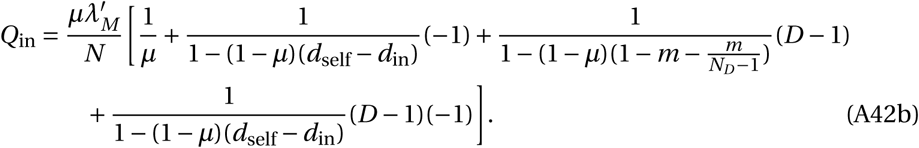

And when *r*_1_ ≢ 0, the two individuals are in different demes:

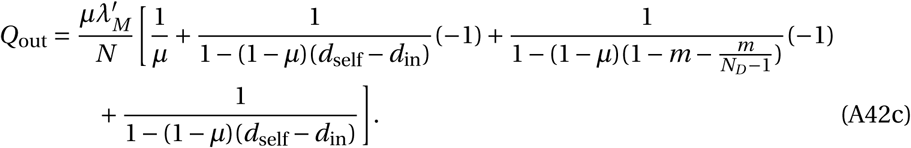

With *d*_self_ *= d*_in_ *=* (1 −*m*)/*n*, we eventually obtain:

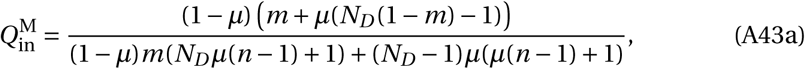

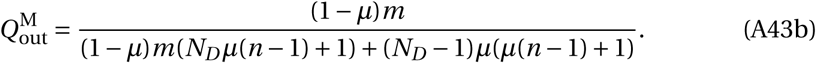

The probability that two different deme-mates are identical by descent, 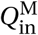, decreases monotonically with the emigration probability *m*, while 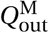 monotonically increases with *m* (see figure A5 (a)).

When the mutation probability *µ* is vanishingly small (*µ* → 0), both 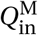 and 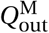 are equal to 1: in the absence of mutation indeed, the population ends up fixed for one of the two types, and all individuals are identical by descent. Note that we obtain a different result if we first assumed that the size of the population is infinite (*N*_*D*_ → *∞*), because the order of limits matters; for instance, 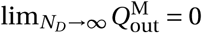.

Relatedness *R* was defined in eq. (A20) as

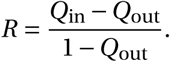

Using eq. (A43), relatedness under the Moran model is given by

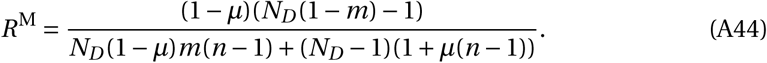

When there is an infinite number of demes (*N*_*D*_ → *∞*) and mutation is vanishingly small (*µ* → 0), we recover

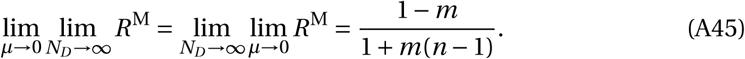

##### C.2.2 Wright-Fisher

For the Wright-Fisher updating, the equation for 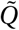 is different:

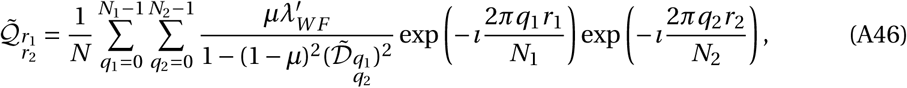

with 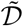 given in eq. (A38b). In a subdivided population, with *N*_1_ *= N*_*D*_ and *N*_2_ *= n*, this becomes

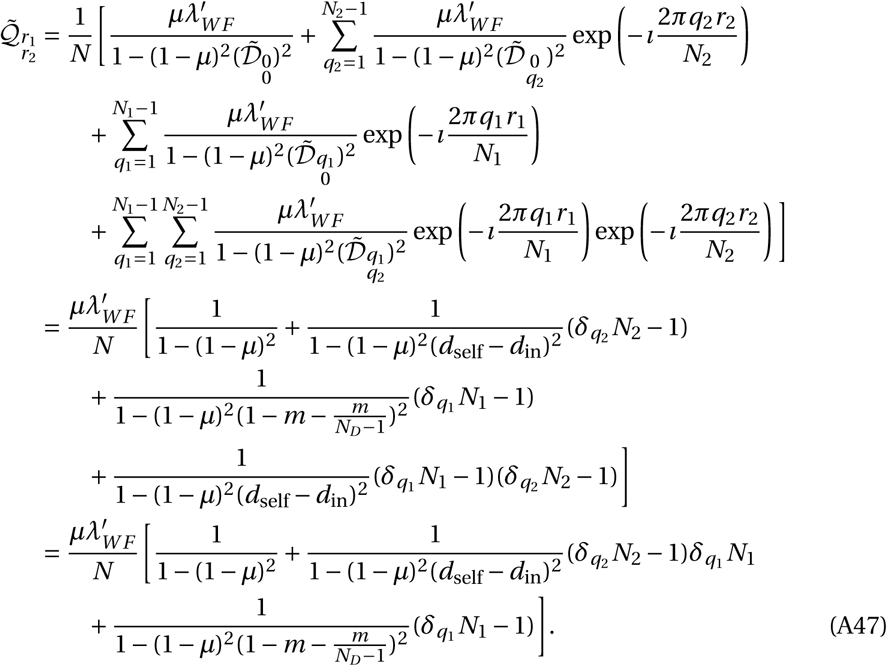

To find 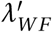, we solve 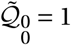, *i.e.*,

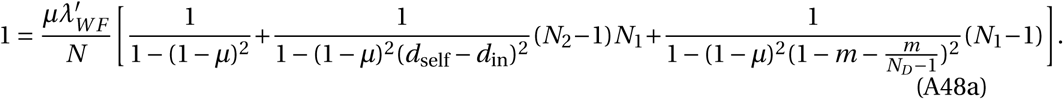

Then from eq. (A47) we deduce

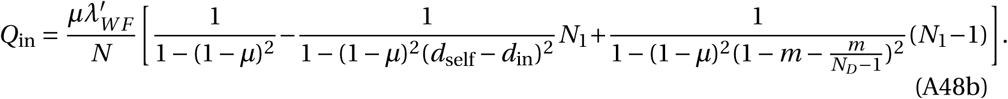

and

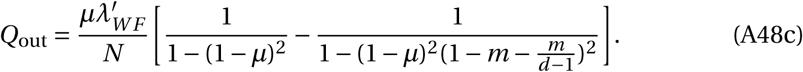

With *d*_self_ *= d*_in_ *=* (1 −*m*)/*n*, we obtain:

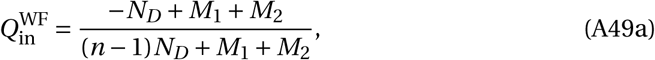

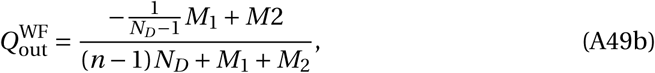

with

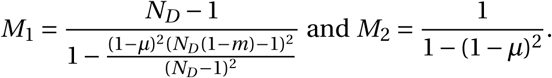

(These formulas are compatible with, *e.g.*, results presented by Cockerham & Weir (1987), adapted for haploid individuals).

In the Wright-Fisher life cycle, 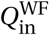 decreases until 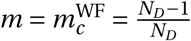, while 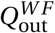follows the opposite pattern. The threshold value 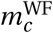 corresponds to an emigration probability so high that *d*_in_ *= d*_out_.

The two probabilities of identity by descent go to 1 when the mutation probability *µ* is very small (*µ* → 0), except if we first assume that the number of demes is very large (*N*_*D*_ → *∞*); for instance, with this life cycle as well, 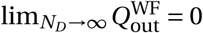.

Also, because more sites (all of them, actually) are updated at each time step, *Q*_in_ is lower for the Wright-Fisher updating than for a Moran updating, under which only one site is updated at each time step (compare figure A5 (a) and A5(b)).

**Figure A5:**
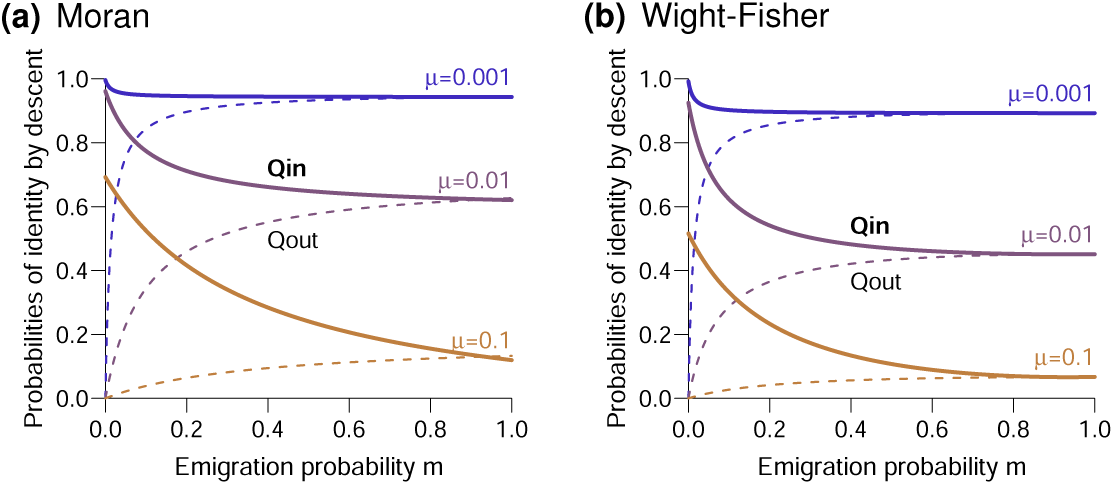
Probabilities of identity by descent, for two different individuals within the same deme (*Q*_in_, full curves) and two individuals in different demes (*Q*_out_, dashed curves), as a function of the emigration probability *m*, for different values of the mutation probability *µ* (0.001, 0.01, 0.1), and for the two types of life cycles ((a): Moran, (b): Wright-Fisher). Other parameters: *n =* 4 individuals per deme, *N*_*D*_ *=* 15 demes.

Combining the formulas presented in eq. (A49), we obtain

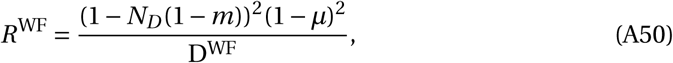

with

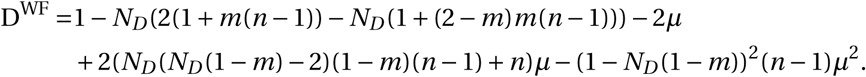

When the number of demes is very large and mutation is vanishingly small, eq. (A50) reduces to

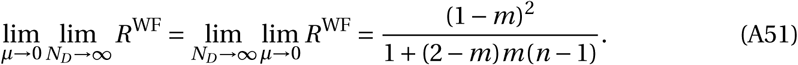

## Footnotes

1 Note that for the sake of concision, we use the word “mutation” throughout the paper, keeping in mind that strategy transmission does not have to be genetic.

## Literature Cited

Alizon, S. & Taylor, P. 2008: Empty sites can promote altruistic behavior. Evolution 62(6):1335–1344.

Allen, B.; Lippner, G.; Chen, Y.-T.; Fotouhi, B.; Momeni, N.; Yau, S.-T. & Nowak, M. A. 2017: Evolutionary dynamics on any population structure. Nature 544(7649):227–230.

Allen, B. & Nowak, M. A. 2014: Games on graphs. EMS surveys in mathematical sciences 1(1):113–151.

Allen, B. & Tarnita, C. E. 2014: Measures of success in a class of evolutionary models with fixed population size and structure. Journal of Mathematical Biology 68(1):109–143.

Allen, B.; Traulsen, A.; Tarnita, C. E. & Nowak, M. A. 2012: How mutation affects evolutionary games on graphs. Journal of Theoretical Biology 299:97–105. Evolution of Cooperation.

Boyd, R. & Richerson, P. J. 2002: Group beneficial norms can spread rapidly in a structured population. Journal of theoretical biology 215(3):287–296.

Brockhurst, M. A.; Buckling, A. & Gardner, A. 2007: Cooperation peaks at intermediate disturbance. Current Biology 17(9):761–765.

Chen, X.; Åke Brännström & Dieckmann, U. 2019: Parent-preferred dispersal promotes cooperation in structured populations. Proceedings of the Royal Society B: Biological Sciences 286(1895):20181949.

Cockerham, C. C. & Weir, B. 1987: Correlations, descent measures: drift with migration and mutation. Proceedings of the National Academy of Sciences 84(23):8512–8514.

Débarre, F. 2017: Fidelity of parent-offspring transmission and the evolution of social behavior in structured populations. Journal of Theoretical Biology 420:26–35.

Débarre, F.; Hauert, C. & Doebeli, M. 2014: Social evolution in structured populations. Nature Communications 5.

Fletcher, J. A. & Doebeli, M. 2009: A simple and general explanation for the evolution of altruism. Proceedings of the Royal Society B: Biological Sciences 276(1654):13–19.

Frank, S. A. 1997: The Price equation, Fisher’s fundamental theorem, kin selection, and causal analysis. Evolution 51(6):1712–1729.

Frank, S. A. 2010: Microbial secretor–cheater dynamics. Philosophical Transactions of the Royal Society of London B: Biological Sciences 365(1552):2515–2522.

Grafen, A. & Archetti, M. 2008: Natural selection of altruism in inelastic viscous homogeneous populations. Journal of Theoretical Biology 252(4):694–710.

Hamilton, W. 1964: The genetical evolution of social behaviour. i. Journal of Theoretical Biology 7(1):1–16.

Hamilton, W. D. 1975: Innate social aptitudes of man: an approach from evolutionary genetics. Biosocial anthropology 53:133–55.

Hammerschmidt, K.; Rose, C. J.; Kerr, B. & Rainey, P. B. 2014: Life cycles, fitness decoupling and the evolution of multicellularity. Nature 515(7525):75–79.

Kandori, M.; Mailath, G. J. & Rob, R. 1993: Learning, mutation, and long run equilibria in games. Econometrica 61(1):29–56.

Le Galliard, J.; Ferrière, R. & Dieckmann, U. 2005: Adaptive evolution of social traits: Origin, trajectories, and correlations of altruism and mobility. The American Naturalist 165(2):206–224. PMID: 15729651.

Lehmann, L.; Feldman, M. W. & Foster, K. R. 2008: Cultural transmission can inhibit the evolution of altruistic helping. The American Naturalist 172(1):12–24. PMID: 18500938.

Lehmann, L.; Keller, L. & Sumpter, D. J. T. 2007: The evolution of helping and harming on graphs: the return of the inclusive fitness effect. Journal of Evolutionary Biology 20(6):2284–2295.

Lehmann, L. & Rousset, F. 2014: The genetical theory of social behaviour. Philosophical Transactions of the Royal Society of London B: Biological Sciences 369(1642).

Leturque, H. & Rousset, F. 2002: Dispersal, kin competition, and the ideal free distribution in a spatially heterogeneous population. Theoretical Population Biology 62(2):169–180.

Lion, S. 2016: Moment equations in spatial evolutionary ecology. Journal of theoretical biology 405:46–57.

Mullon, C.; Keller, L. & Lehmann, L. 2017: Co-evolution of dispersal with behaviour favours social polymorphism. bioRxiv.

Mullon, C. & Lehmann, L. 2014: The robustness of the weak selection approximation for the evolution of altruism against strong selection. Journal of evolutionary biology 27(10):2272–2282.

Ohtsuki, H.; Hauert, C.; Lieberman, E. & Nowak, M. A. 2006: A simple rule for the evolution of cooperation on graphs and social networks. Nature 441(7092):502–505.

Parvinen, K. 2013: Joint evolution of altruistic cooperation and dispersal in a metapopulation of small local populations. Theoretical population biology 85:12–19.

Platt, T. G. & Bever, J. D. 2009: Kin competition and the evolution of cooperation. Trends in Ecology & Evolution 24(7):370–377.

Rodrigues, A. M. M. & Gardner, A. 2012: Evolution of helping and harming in heterogeneous populations. Evolution 66(7):2065–2079.

Rousset, F. 2004: Genetic Structure and Selection in Subdivided Populations. Princeton University Press, Princeton, NJ.

Rousset, F. & Billiard, S. 2000: A theoretical basis for measures of kin selection in subdivided populations: finite populations and localized dispersal. Journal of Evolutionary Biology 13(5):814–825.

Sample, C. & Allen, B. 2017: The limits of weak selection and large population size in evolutionary game theory. Journal of mathematical biology pages 1–33.

Schneider, D. M.; Martins, A. B. & de Aguiar, M. A. 2016: The mutation–drift balance in spatially structured populations. Journal of theoretical biology 402:9–17.

Tarnita, C. E. & Taylor, P. D. 2014: Measures of relative fitness of social behaviors in finite structured population models. The American Naturalist 184(4):477–488.

Taylor, P. 1992a: Altruism in viscous populations—an inclusive fitness model. Evolutionary ecology 6(4):352–356.

Taylor, P. 2010: Birth–death symmetry in the evolution of a social trait. Journal of Evolutionary Biology 23(12):2569–2578.

Taylor, P.; Lillicrap, T. & Cownden, D. 2011: Inclusive fitness analysis on mathematical groups. Evolution 65(3):849–859.

Taylor, P. D. 1992b: Inclusive fitness in a homogeneous environment. Proceedings of the Royal Society of London. Series B: Biological Sciences 249(1326):299–302.

Taylor, P. D.; Day, T. & Wild, G. 2007a: Evolution of cooperation in a finite homogeneous graph. Nature 447(7143):469–472.

Taylor, P. D.; Day, T. & Wild, G. 2007b: From inclusive fitness to fixation probability in homogeneous structured populations. Journal of Theoretical Biology 249(1):101–110.

Taylor, P. D. & Irwin, A. J. 2000: Overlapping generations can promote altruistic behavior. Evolution 54(4):1135–1141.

Traulsen, A.; Hauert, C.; De Silva, H.; Nowak, M. A. & Sigmund, K. 2009: Exploration dynamics in evolutionary games. Proceedings of the National Academy of Sciences 106(3):709–712.

Van Cleve, J. 2015: Social evolution and genetic interactions in the short and long term. Theoretical Population Biology 103:2–26.

West, S. A. & Gardner, A. 2010: Altruism, spite, and greenbeards. Science 327(5971):1341–1344.

West, S. A.; Pen, I. & Griffin, A. S. 2002: Cooperation and competition between relatives. Science 296(5565):72–75.

Wild, G. & Traulsen, A. 2007: The different limits of weak selection and the evolutionary dynamics of finite populations. Journal of Theoretical Biology 247(2):382–390.

Wilson, D. S.; Pollock, G. B. & Dugatkin, L. A. 1992: Can altruism evolve in purely viscous populations? Evolutionary Ecology 6(4):331–341.

Wolfram Research, Inc. 2017: Mathematica, Version 11.1. Champaign, IL, 2017.

